# Hypoxia-induced CTCF mediates alternative splicing via coupling chromatin looping and RNA Pol II pause to promote EMT in breast cancer

**DOI:** 10.1101/2023.05.06.539689

**Authors:** Parik Kakani, Shruti Dhamdhere, Deepak Pant, Jharna Mishra, Atul Samaiya, Sanjeev Shukla

## Abstract

Cancer cells experiencing hypoxic stress employ Epithelial-Mesenchymal Transition (EMT) to undergo metastasis through rewiring chromatin landscape, epigenetics, and importantly alternative splicing. Here, we investigated the role of CTCF, chromatin, epigenetic and alternative splicing modulator under hypoxia to promote EMT. Our result shows that hypoxia-induced epigenetic changes upregulate *CTCF* expression in breast cancer. We delineate that *CTCF* is a direct target of HIF1α and the hypoxia-induced HIF1α-CTCF axis functionally contributes to promoting EMT in breast cancer cells. Finally, our work uncovers *COL5A1*, an EMT gene, as a direct target of CTCF. We demonstrated that hypoxia-mediated CTCF enrichment on *COL5A1* promoter regulates expression as well as alternative splicing events to promote EMT. Here, we put forward an intricate mechanism of alternative splicing where CTCF-mediated promoter-exon upstream looping regulates DNA de-methylation and CTCF-mediated RNA Pol II pausing at *COL5A1* exon 64A. Our global analysis of CTCF-ChIP-seq data reveals that the hypoxia-induced differential CTCF occupancy possibly regulates gene expression and alternative splicing events of many genes that are enriched in the EMT pathway, cancer cell motility and invasion, angiogenesis, and stemness under hypoxia, similarly to the proposed model. Finally, we employed the epigenetic modulator dCas9-DNMT3A system to specifically disrupt HIF1α or CTCF binding under hypoxia and hence the HIF1α-CTCF-COL5A1exon64A axis that alleviates the EMT potential of breast cancer cells that may represent a novel therapeutic target in breast cancer.

**Graphical Abstract.**
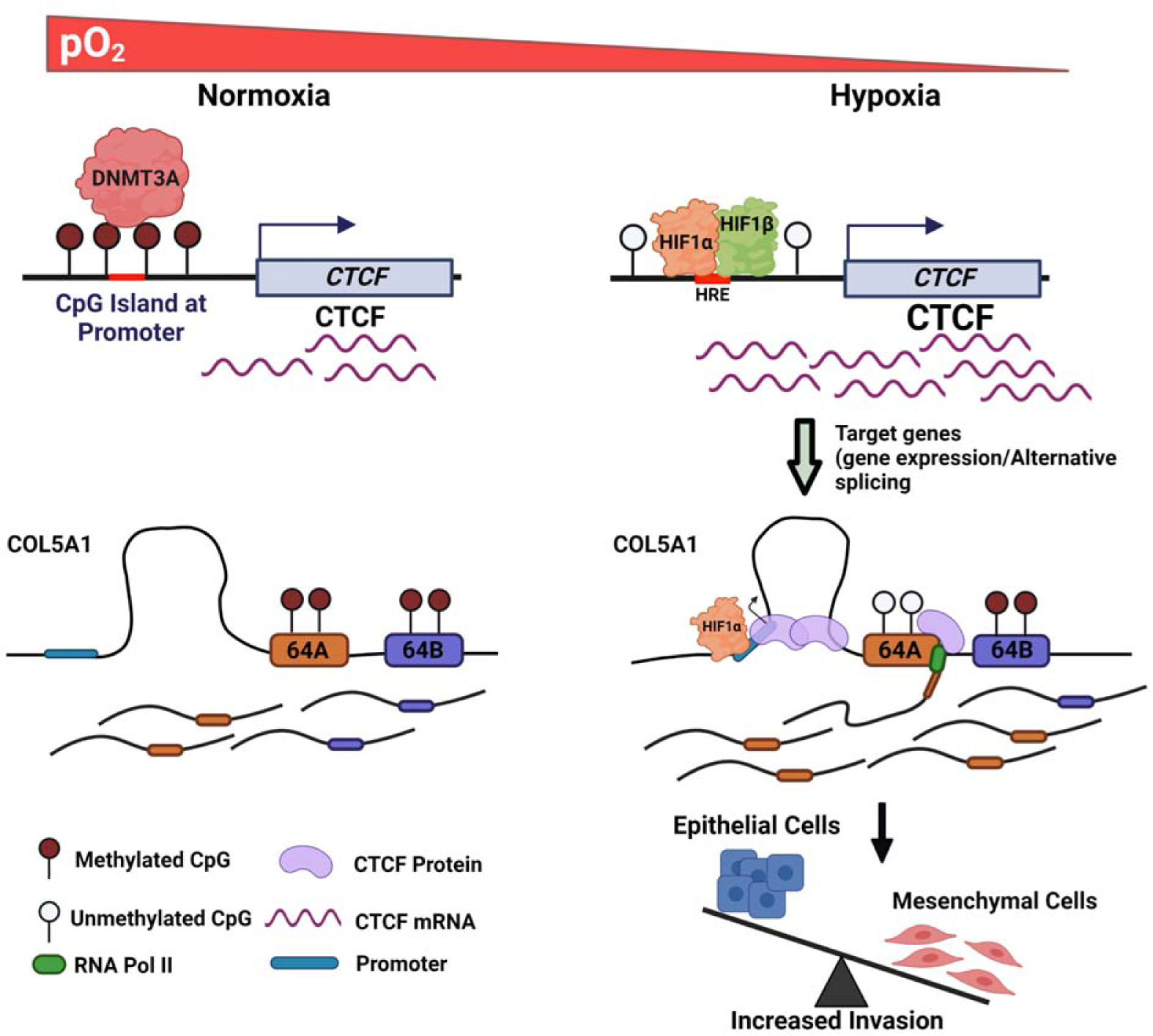

## Introduction

Cancer being a heterogenous and complex pathological condition, contributes heavily to the overall world morbidity and mortality. The ability of cancer cells to traverse through the vasculature to reach and proliferate in the surrounding tissues as well as distant organs, termed as metastasis, is the primary cause of poor prognosis and fatality ^1^. Hypoxia, a low oxygen stress condition present in the core of solid tumors, is associated with cancer progression, invasion, angiogenesis, changes in metabolism, and increased metastasis ^2^. Conclusively, hypoxia is a potent micro-environmental factor that promotes tumor metastasis ^1–3^. Extensive changes in DNA methylation landscape, epigenetic as well as gene expression reprogramming, and most importantly, alternative splicing have been characterized as a survival strategies adopted by cancer cells to escape and overcome from hypoxic stress, which also renders a more aggressive phenotype ^4–12^. Moreover, a plethora of studies have reported the regulatory role of epigenetic modifications in differential gene expression and alternative splicing of oncogenic genes that drive tumor initiation and early progression ^13–18^. CCCTC-binding factor (CTCF), a multifunctional DNA binding protein, is essentially involved in regulating epigenetic reprogramming, gene expression, and alternative splicing ^19–21^. Interestingly, despite the involvement of CTCF in epigenetic reprogramming and alternative splicing, there are yet no reports discussing the role of CTCF in shaping hypoxic cancer transcriptome.

In this study, we demonstrate that hypoxia induces *CTCF* expression via cumulative effect of DNA methylation and recruitment of HIF1α on *CTCF* promoter. Furthermore, we find that the hypoxia-driven HIF1α-CTCF axis regulates the EMT process and decides invasive properties of cancer cells. Here, we showed hypoxia-driven CTCF-dependent tripartite relationship between epigenetic reprogramming, alternative splicing, and EMT in the cancer cell. Our ChIP-seq analysis showed hypoxia-driven differential CTCF binding sites overlapped with many EMT and cancer pathways related genes and possibly regulate their expression as well as alternative splicing. Here, we identified *COL5A1* a hypoxia-induced EMT gene, as a novel target of CTCF and hypoxia-induced CTCF regulates *COL5A1* expression as well as alternative splicing to promote invasive phenotype. We demonstrate that hypoxia promotes exon 64A inclusion in *COL5A1* mRNA that favors EMT. The inclusion of exon 64A is regulated by CTCF-mediated intricate mechanism coupling CTCF-mediated promoter-exon upstream looping with CTCF-mediated RNA Pol II pausing at exon 64A. Finally, we use epigenome editing strategy using dCas9-DNMT3A system for potent, specific, and stable disruption of HIF1α or CTCF binding and hence HIF1α-CTCF-COL5A1exon 64A axis that alleviates EMT potential of breast cancer cells under hypoxia. Thus, our data demonstrate a specific function of the chromatin organizer CTCF for hypoxia-driven EMT by modulating alternative splicing and chromatin organization.

## Results

### *CTCF* is induced in hypoxic breast cancer

To apprehend the hypoxia driven differential expression of *CTCF*, breast cancer cell lines MCF7 and HCC1806 were cultured under normoxic (21% O_2_) or hypoxic (1% O_2_) conditions. We analyzed the effect of hypoxia on CTCF protein expression by immunoblot assay and found that CTCF protein levels were significantly increased, along with HIF-1α protein, under hypoxic condition **(Figure 1A and 1B)**. To provide direct evidence for hypoxia-induced transcriptional upregulation of *CTCF* expression in breast cancer cell lines, we checked the mRNA levels of *CTCF* using qPCR. Data presented in **Figure 1C and 1D** showed increased mRNA levels of *CTCF* in both MCF7 and HCC1806 breast cancer cell lines under hypoxia as compared to normoxic cells. These observations indicated that transcriptional upregulation of *CTCF* expression is accounting for its increased protein expression under hypoxia.

**Figure 1:**
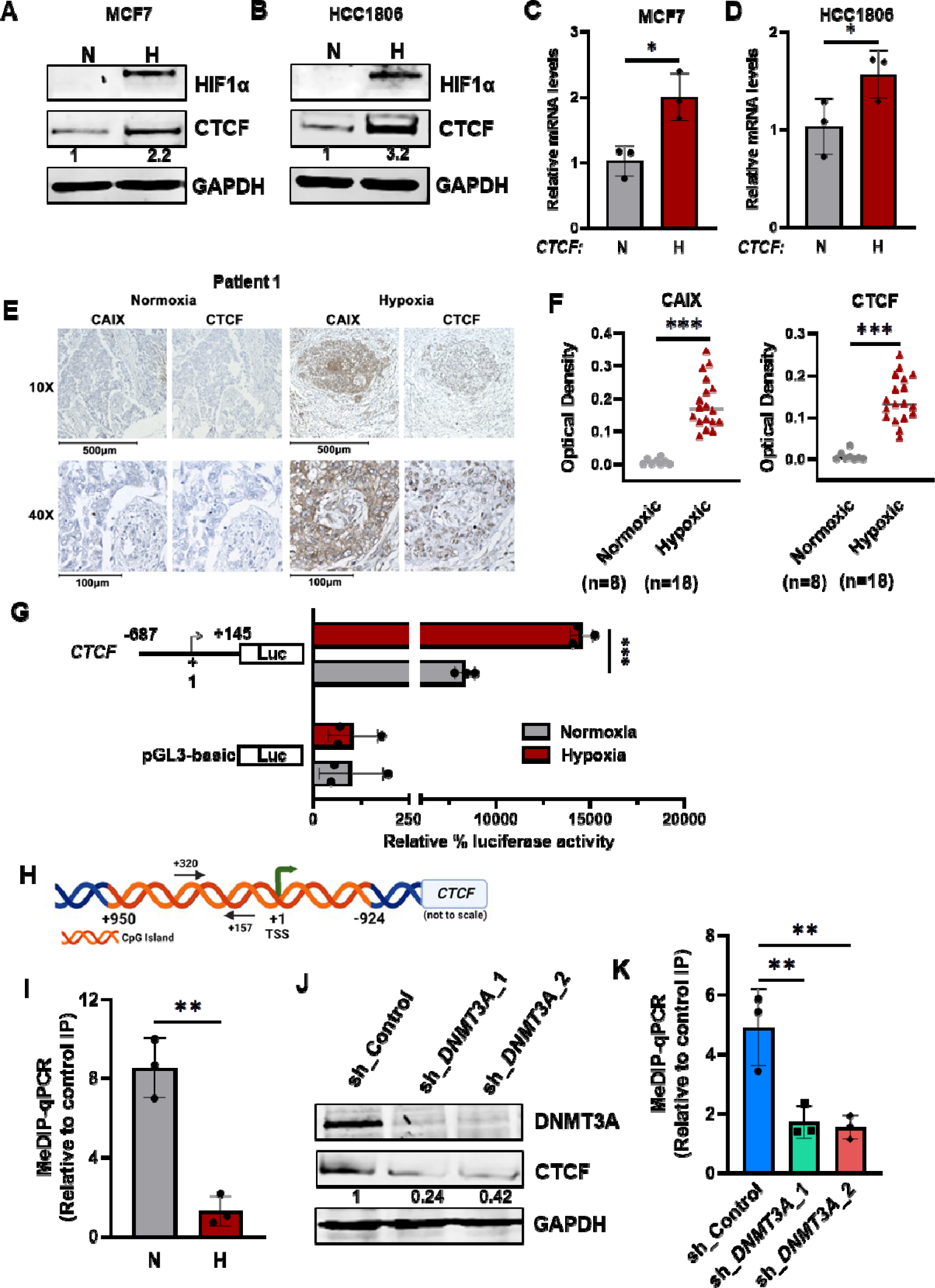
CTCF is induced in hypoxic breast cancer. **A) and B)** Immno-blot of CTCF and HIF1α in normoxic and hypoxic breast cancer cell line, MCF7 and HCC1806, GAPDH is used as loading control here (and elsewhere) **C) and D)** qRT-PCR of CTCF in normoxic and hypoxic breast cancer cell line, MCF7 and HCC1806 normalized to RPS16 mRNA levels **E)** CAIX and CTCF immunostaining of illustrative case of breast cancer patients **F)** and its quantification (n=18) represented as optical density for hypoxic versus normoxic regions of breast cancer patients, **G)** MCF7 cells were co-transfected with CTCF promoter-luciferase construct (−687 to +145) and pRL-TK Renilla, exposed to normoxia and hypoxia. The relative luciferase values are shown as mean ± SD **H)** schematic showing presence of dense CpG island in CTCF promoter around TSS **I)** MeDIP-qPCR on CTCF promoter in normoxic vs hypoxic MCF7 cells **J)** Immunoblot of DNMT3A and CTCF in normoxic MCF7 cells transduced with shRNA against DNMT3A or shControl **K)** MeDIP-qPCR on CTCF promoter in normoxic MCF7 cells transduced with shRNA against DNMT3A versus shControl. Error bar shows mean values ± SD. (n=3). As calculated using two-tailed Student’s t test, *p < 0.05, **p < 0.01, ***p < 0.001.

To determine the clinical significance of hypoxia-induced CTCF expression in breast cancer patients, we investigated the association between hypoxic region and CTCF expression. We performed immunohistology analyses of breast tissue sections obtained from breast cancer patients. The carbonic anhydrase IX (CAIX), which is previously reported as a biomarker of hypoxia ^9, 22^, was used to immunostain the sections in parallel with CTCF. The regions corresponding to positive CAIX staining showed strong CTCF expression **(Figure 1E and Figure S1)**. Moreover, normoxic regions (negatively stained for CAIX) of the tumor tissue correspond to weak CTCF expression. The DAB staining when quantified for all 18 patients shows strong signal for CAIX and CTCF in hypoxic region compared to normoxic regions (even in the same patient), (**Figure 1F**). These results corroborate with our observations in the breast cancer cell lines that hypoxic tumor cells are associated with increased *CTCF* expression. Thus, strengthen our finding that hypoxia induced *CTCF* expression in breast cancer.

We next expanded our study and test whether CTCF expression is correlated with the expression of hypoxia metagene signature (195) consisting of HIF-regulated genes that play key roles in breast cancer metastasis such as *CXCR3*, *L1CAM*, *LOX*, *P4HA1*, *P4HA2*, *PDGFB*, *PLOD1*, *PLOD2*, *SLC2A1*, and *VEGFA* (list from MSigDB). Using GEPIA 2.0 platform, we did Pearson’s test and found a significant correlation between *CTCF* mRNA levels with the hypoxia signature genes mRNA levels with a positive correlation coefficient of R = 0.35 (**Figure S2A**). Additionally, analysis of mRNA expression data from 1,218 breast tissue specimens in the Cancer Genome Atlas (TCGA) database showed a non-significant change in *CTCF* expression in normal vs tumor breast tissue **(Figure S2B)**. However, when TCGA 1090 tumor samples were stratified as CTCF_high or CTCF_low showed enrichment of hypoxia hall mark geneset (SHI_ET_AL_2007) in CTCF_high tumor patients as compared to CTCF_low tumor patients (**Figure S2C**).

### Hypoxia-driven differential DNA methylation at *CTCF* promoter regulates its expression and is maintained by DNMT3A under normoxia in breast cancer cell lines

We then investigated the molecular mechanisms involved in the upregulation of *CTCF* expression under hypoxia. We did a dual luciferase reporter assay to investigate the promoter sequence involved in hypoxia-mediated transcriptional upregulation of *CTCF*. For that 5′-upstream sequence of *CTCF* transcription start site (TSS) was amplified and cloned into pGL3-basic vector, pGL3-CTCF (−687 to +145 bp) (**Figure 1G**). Dual luciferase assay revealed that the luciferase activity of pGL3-*CTCF* promoter construct was significantly increased in hypoxic condition as compared to the normoxia in breast cancer cell lines **(Figure 1G and Figure S2D)**, suggesting the presence of potential positive regulatory elements in this region, which enhances *CTCF* transcription under hypoxia.

To identify the transcription factor (TF) and molecular mechanism underlying the hypoxia-driven *CTCF* induction, we further analyzed the *CTCF* promoter region around TSS to mark the presence of regulatory elements. Interestingly, promoter analysis revealed the presence of dense CpG Island in the promoter region of *CTCF* spanning TSS (**Figure 1H)**. Several studies reported the role of DNA methylation around CpG Island to drive the temporal-spatial expression of genes ^23^. We checked the methylation levels of *CTCF* promoter around CpG Island in breast cancer cell lines, and MeDIP-qPCR revealed methylated *CTCF* promoter around TSS under normoxia (**Figure 1I and Figure S2E**).

Based on the above findings, we next sought to understand the involvement of DNA methyltransferases (s) (DNMT1, DNMT3A, and DNMT3B) in maintaining DNA methylation at *CTCF* promoter under normoxic condition. We next asked whether modulating the expression of *DNMT1* or *DNMT3A* or *DNMT3B* affects methylation status of *CTCF* promoter, we knockdown all the three DNMTs individually (**Figure 1J and Figure S2F and S2G**) and checked the CTCF promoter methylation status using MeDIP-qPCR in these normoxic breast cancer cells. Data presented in **Figure S2I and S2J** revealed that depletion of either DNMT1 or DNMT3B had no effect on DNA methylation at *CTCF* promoter but significantly lowered the amount of DNA methylation at *CTCF* promoter when DNMT3A was depleted (**Figure 1J and 1K and Figure S2H and S2K**). These results confirmed that DNMT3A is the one responsible for maintaining the DNA methylation around TSS at *CTCF* promoter. Further, we checked the expression of CTCF upon DNMTs KD in normoxic breast cancer cells, and we found that knockdown of both DNMT1 and DNMT3B did not show any significant change in CTCF expression (**Figure S2F and S2G**) but surprisingly, DNMT3A knockdown greatly reduced CTCF expression in normoxic cells (**Figure 1J and Figure S2H**). This implies that, indeed DNA methylation around TSS on CTCF promoter plays a significant role and is positively correlated with *CTCF* expression. In fact, very few studies have reported positive role of DNA methylation in gene expression ^24^. Collectively, these data demonstrate that DNMT3A-mediated DNA methylation at *CTCF* promoter is a major mechanism responsible for maintaining *CTCF* expression in breast cancer cells. These observations warrant further studies to define what type of protein, a transcriptional activator that has preferred binding to methylated DNA or transcriptional repressor unable to bind due to methylated DNA is regulating *CTCF* expression.

Further, we checked methylation levels in hypoxic conditions to explore the role of DNA methylation at CpG Island in regulating the expression of *CTCF* under hypoxic condition. Surprisingly, we observed reduced methylation at *CTCF* promoter around TSS in hypoxia as compared to normoxia (**Figure 1I and Figure S2D)**. Altogether, these data encouraged us to hypothesize that CTCF expression is regulated by an epigenetic mechanism that includes differential DNA methylation at CTCF promoter under hypoxic condition and suggested the involvement of hypoxia-specific transcription activator in mediating CTCF upregulation. We further expanded our study to define the precise mechanism(s) regulating hypoxia-induced CTCF expression and to decipher their relative contributions in the acquisition of hypoxia-mediated aggressive phenotypes in breast cancer.

### Hypoxia-induced CTCF expression is HIF1**α**-dependent

To identify the transcriptional regulator responsible for the upregulation of *CTCF* expression in hypoxia, we analyzed its promoter region for transcription factor binding motifs. Interestingly, we found putative binding motifs for hypoxia-specific transcriptional activator, HIF1α, known as hypoxia-responsive elements (HREs) at +262 bp and +248 bp upstream of the *CTCF* TSS **(Figure 2A)**. To validate the *in silico* findings, we performed HIF1α ChIP-qPCR on *CTCF* promoter in normoxic vs. hypoxic condition, and data presented in **Figure 2B and Figure S3A** showed HIF1α enrichment on *CTCF* promoter under hypoxia.

**Figure 2:**
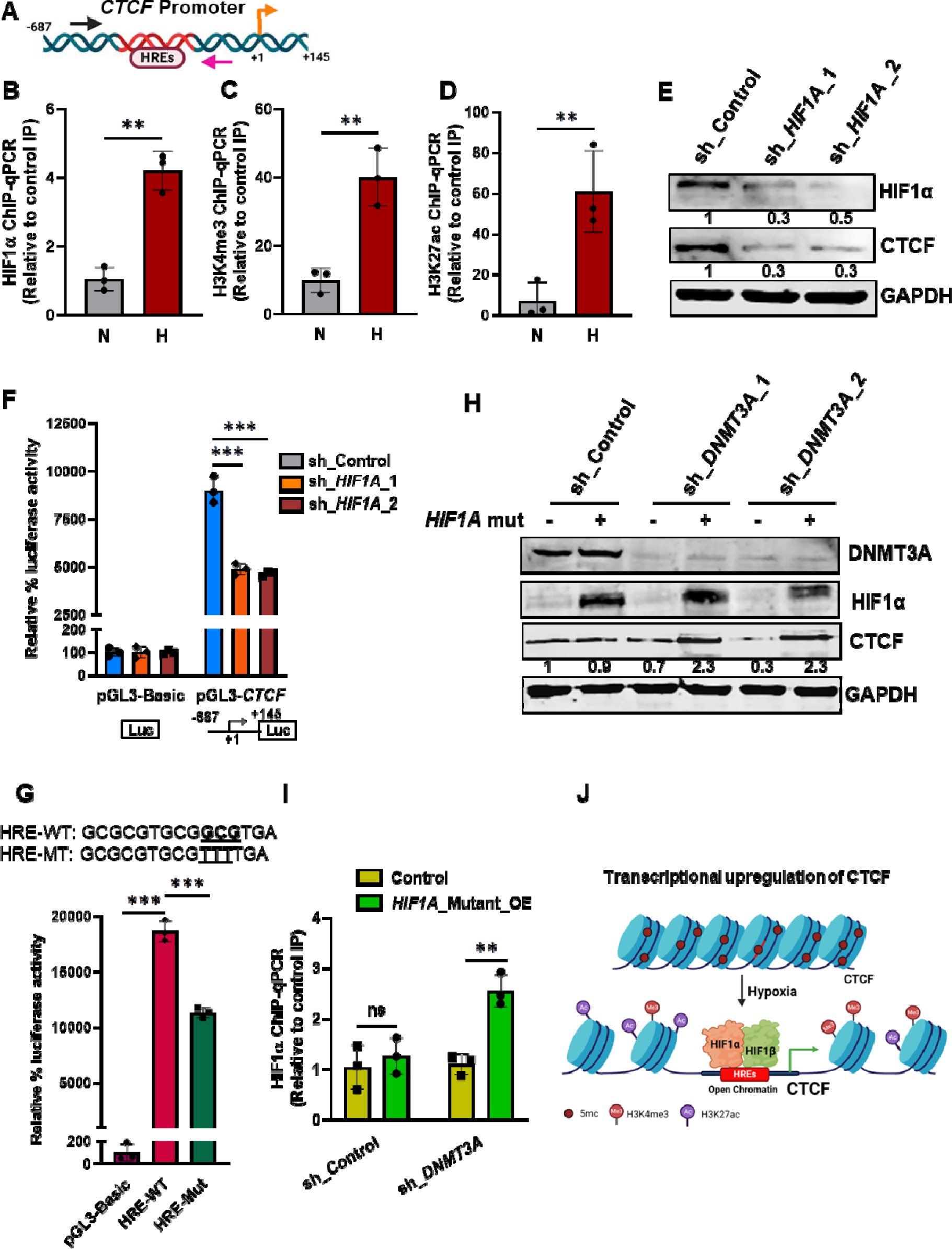
CTCF is a direct target of HIF1α. **A)** Schematic showing presence of putative HRE elements on CTCF promoter **B)** HIF1α ChIP-qPCR **C)** H3K4me3 ChIP-qPCR **D)** H3K27ac ChIP-qPCR on CTCF promoter in normoxic and hypoxic MCF7 cells **E)** Immunoblot of CTCF after HIF1α knockdown in MCF7 cells under hypoxia **F)** Relative luciferase activity of CTCF promoter in MCF7 cells transfected with shRNA against HIF1α vs shcontrol under hypoxia **G)** Schematic representation of the HRE sequences for both wild-type and mutated constructs and wild-type or mutant CTCF-promoter luciferase constructs were co-transfected with the Renilla luciferase vector in MCF7 cells under hypoxia, and the luciferase activity was measured **H)** Immunoblot of CTCF, DNMT3A and HIF1α after HIF1A mutant (Addgene 19005) overexpression in DNMT3A depleted cells or control in MCF7 under normoxia **I)** HIF1α ChIP-qPCR on CTCF promoter after HIF1A mutant (Addgene 19005) overexpression in DNMT3A depleted cells or control in MCF7 under normoxia **J)** Schematic showing hypoxia-driven epigenetic and transcriptional regulation of CTCF expression in breast cancer cell. Error bar shows mean values ± SD. (n=3). As calculated using two-tailed Student’s t test, *p < 0.05, **p < 0.01, ***p < 0.001.

Hypoxia causes increased H3K4me3 and H3K27ac marks at HIF1α target genes in HIF1α dependent manner ^25^. We measured the levels of H3K4me3 and H3K27ac marks at *CTCF* promoter around HRE region to validate hypoxia-specific upregulation of *CTCF* and as a direct transcriptional target of HIF1α. Results presented in **Figure 2C and 2D and Figure S3B and S3C** revealed that hypoxia causes enrichment of both H3K4me3 and H3K27ac marks around HRE consistent with active gene transcription ^25^, providing further evidence for promoting HIF1α-mediated *CTCF* transcription in hypoxia.

Next, to verify the transcriptional modulation of *CTCF* by HIF1α under hypoxia, we checked CTCF expression in HIF1α depleted cells under hypoxia. It was observed that sh-RNA mediated knockdown of *HIF1A* in MCF7 significantly reduced CTCF expression compared to the shcontrol under hypoxia **(Figure 2E)**. Similarly, CTCF expression was also assessed in *HIF1A* knock out (KO) HCC1806 cells under hypoxic condition. Data presented in **Figure S3D** revealed that a complete knock out of HIF1α led to reduced expression of CTCF under hypoxia. Moreover, luciferase activity of pGL3-*CTCF*-promoter luciferase construct decreased drastically in both *HIF1A* KD MCF7 and *HIF1A* KO HCC1806 cells compared to the control cells under hypoxia (**Figure 2F and Figure S3E)**. Next, we investigated whether DNA fragments encompassing HIFα binding sites function as hypoxia response elements (HREs). We mutated HRE present in the *CTCF*-promoter luciferase construct and performed luciferase assay. Exposure to hypoxia resulted in a significant increase in the wild-type pGL3-*CTCF*-promoter-luciferase construct, whereas mutation of the HIF1α binding sequence within HREs decreased hypoxia-induced luciferase activity (**Figure 2G**), demonstrating that 5’-GGCGTGA-3’ functions as an HREs. Taken together, we demonstrate that HIF-1α binds directly to the *CTCF* gene promoter to activate transcription in hypoxic breast cancer cells.

We further set out to investigate whether hypoxia induced DNA de-methylation at *CTCF* promoter is HIF1α dependent or independent. To answer this, we overexpressed *HIF1A* mutant plasmid (**Figure S3F**) that produced stable form of HIF1α protein under the normoxic condition in control cells (having methylated *CTCF* promoter, **Figure 1I**) or DNMT3A depleted cells (having non methylated *CTCF* promoter, **Figure 1K**). The results presented in **Figure 2H** showed that overexpression of mutant *HIF1A* plasmid produced stable HIF1α protein that resulted in enhanced *CTCF* expression only in DNMT3A depleted cells while no change was observed in normoxic control cells. HIF1α ChIP-qPCR in the same set of cells showed enrichment of HIF1α on *CTCF* promoter only in *HIF1A* mutant plasmid over expressed DNMT3A depleted cells in normoxic condition in comparison to the control cells (**Figure 2I**). Altogether, we conclude that hypoxia-mediated DNA de-methylation at CTCF promoter is HIF1α independent and HIF1α cannot bind methylated DNA consistent with the previous finding that explore the negative correlation between DNA methylation and HIF1α binding in the genome ^26^. Hence, hypoxia-induced epigenetic changes at CTCF promoter allow HIF1α binding that upregulates CTCF expression (**Figure 2J**).

### Hypoxia-induced HIF1**α**-CTCF axis as a major driver of invasive phenotype

Hypoxia promotes EMT and metastatic phenotypes in human cancer cells ^9, 27, 28^. To determine whether hypoxia-induced *CTCF* expression contributes inthe EMT process, we analyzed the effect of *CTCF* knockdown in MCF7 and HCC1806 cells on hypoxia-driven phenotype and invasive behavior of breast cancer cells. The transwell assay results demonstrated that hypoxia significantly increased EMT phenotype and invasive property of both MCF7 and HCC1806 cells, whereas hypoxia-induced EMT was significantly impaired by *CTCF* knockdown as compared to the shControl cells **(Figure 3A, 3B and Figure S4A, S4B)**. Furthermore, we analyzed the expression of EMT molecules such as transcription factor Snail and phenotypic markers, E-cadherin (E-cad) and vimentin and observed consistent results with invasion assay where *CTCF* knockdown abrogates hypoxia-mediated acquisition of mesenchymal phenotype. We observed downregulation of mesenchymal TF Snail and phenotypic marker vimentin, whereas induced epithelial marker E-cad in *CTCF* knockdown cells compared to the shControl under hypoxia (**Figure 3C and Figure 4SC**). Based on these data, we conclude that hypoxia-driven *CTCF* expression is required for the acquisition of EMT phenotype and hence enhanced invasion by hypoxic breast cancer cells. We have shown that HIF1α loss confers reduced *CTCF* expression under hypoxia (**Figure 2C**). Moreover, *CTCF* promotes EMT under hypoxia (**Figure 3**). So, we hypothesized that hypoxia induces HIF1α-CTCF axis, thereby promoting the invasive phenotype of breast cancer cells. To further explore the association, we performed rescue experiment where we overexpressed CTCF in *HIF1A-silenced* MCF7 and HCC1806 cells and the invasive abilities of these cancer cells were assessed by invasion assay.

**Figure 3:**
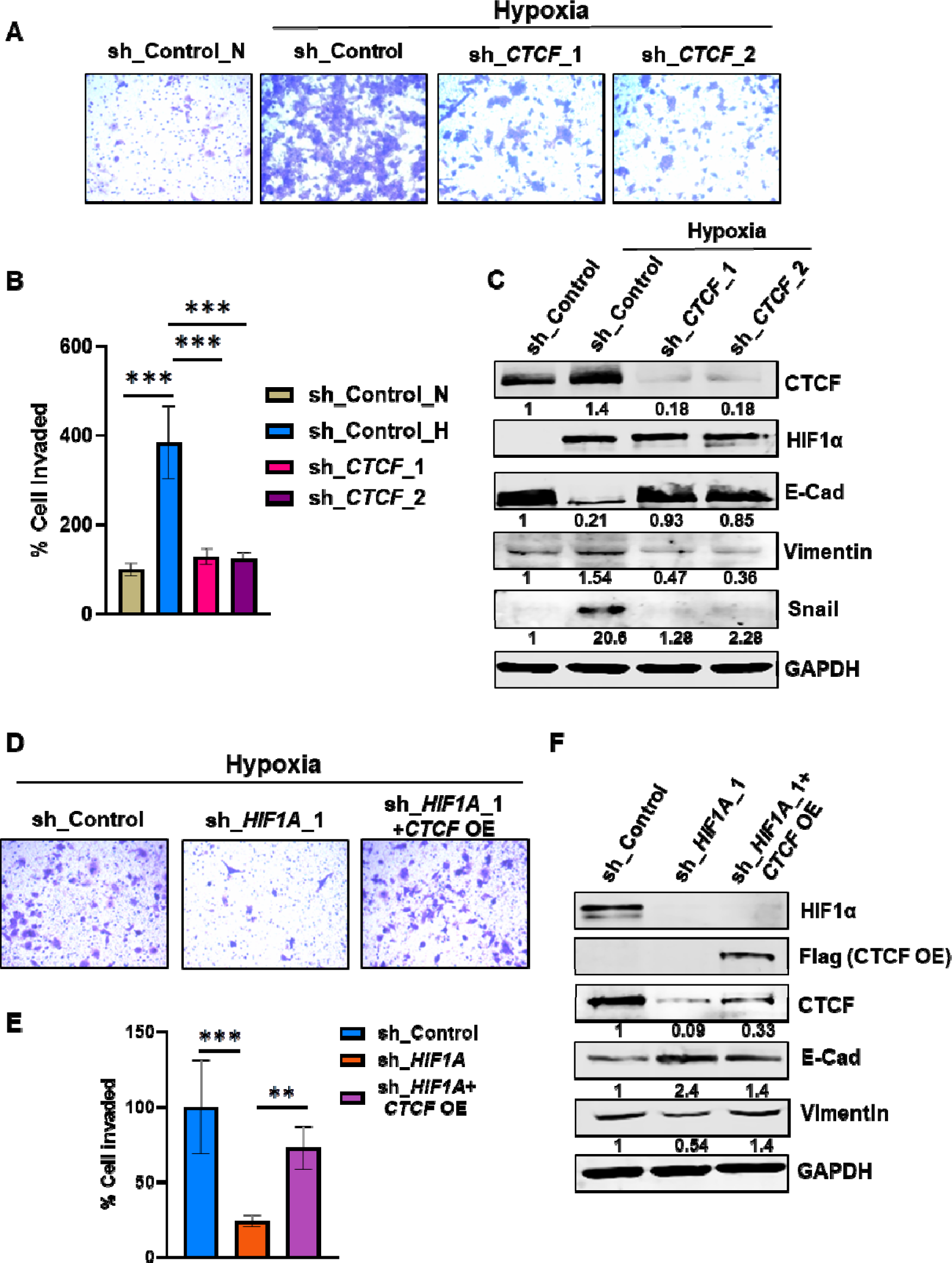
Hypoxia induced HIF1α-CTCF signaling axis promotes breast cancer cell invasive phenotype. **A)** Invasion assay with **B)** its quantification as % of cells invaded after CTCF knockdown in comparison to the control MCF7 cells under normoxic and hypoxic condition **C)** Immunoblot of CTCF, HIF1α, E-cadherin (E-cad), vimentin and snail in CTCF knockdown or control MCF7 cells under normoxic and hypoxic condition **D)** Invasion assay with **E)** quantification as % of cells invaded after overexpression of empty vector (EV) or CTCF in cells transduced with shHIF1A in comparison to the shcontrol under hypoxia in MCF7 **G)** Immunoblot of HIF1α, Flag (to confirm overexpression of CTCF), CTCF, E-cad and vimentin in shControl versus shHIF1A cells with ectopically expressed either empty vector (EV) or CTCF in MCF7 hypoxic cells. Error bar shows mean values ± SD. (n=3). As calculated using two-tailed Student’s t test, *p < 0.05, **p < 0.01, ***p < 0.001.

The reduced HIF1α expression in breast cancer cells resulted in reduced number of cells that have invaded as compared to the shControl under hypoxia (**Figure 3D, 3E and Figure S4D, S4E, S4G, S4H**). When *CTCF* was overexpressed in the *HIF1A* silenced cells, the invasive abilities of hypoxic breast cancer cells were significantly rescued as compared to the *HIF1A* silenced cells under hypoxia (**Figure 3D, 3E and Figure S4D, S4E, S4G, S4H**). To examine the effects of HIF1α-CTCF axis on EMT, western blot analysis was conducted in order to detect the markers of EMT. The expression of E-cad was found to be inhibited, whereas the expression of vimentin was considerably increased after *CTCF* overexpression in *HIF1A-depleted* hypoxic cells (**Figure 3F and Figure S4F and S4I**). These results indicated that CTCF regulates the invasive properties of hypoxic breast cancer cells via the HIF1α signaling pathway, thus promoting the progression of breast cancer.

### Epigenetic modulator dCas9-DNMT3A mediated HREs methylation suppressed hypoxia-induced *CTCF* expression and impaired EMT phenotype

Further, we aimed for targeted epigenetic approach to suppress the *CTCF* expression in hypoxic cancer cell lines. The *CTCF* promoter is de-methylated in hypoxic condition exposing HREs and allow HIF1α to binds and drive activation of *CTCF* expression, which indicates a DNA methylation-dependent molecular control (**Figure 1 and 2**). This prompted us to investigate the direct intrusion with the *CTCF* promoter DNA methylation status to achieve its suppression under hypoxic condition. This would prove to us the therapeutic utility of epigenetically modulated mediated *CTCF* suppression under hypoxia to inhibit cancer invasive phenotype. We used CRISPR-dCas9 (nuclease deficient) fused to epigenetic modulator DNMT3A, which is a DNA methyltransferases ^29^. We designed three overlapping guide RNAs that covered HREs present in the promoter region of *CTCF* responsible for its HIF1α dependent induction under hypoxia (**Figure 4A**). We specifically sought to evaluate the effect of sgRNA-dCas9-DNMT3A system directed to HREs on the *CTCF* expression under hypoxia. To decouple the effect, we did immunoblot assay and found that transient expression of dCas9-DNMT3A along with specific sgRNA reduced CTCF expression under hypoxia in comparison the sgcontrol cells (**Figure 4B**). Our initial result showed that dCas9-DNMT3A guided by sg2 was more effective than other sgRNAs. This indicated that the guide RNA immediately upstream of the HREs is more effective. Hence, sg-RNA2 was further selected for all other analysis. To analyze the effect of dCas9-DNMT3A on DNA methylation status of *CTCF* promoter, we performed MeDIP-qPCR in sgcontrol and dCas9-DNMT3A-sg2 cells under normoxic and hypoxic conditions. We found that dCas9-DNMT3A-sg2 successfully abolishes hypoxia-induced DNA de methylation and maintains methylation levels of the *CTCF* promoter under hypoxia as compared to the sgcontrol cells (**Figure 4C**). Next we asked whether sg2-guided DNMT3A-mediated increased DNA methylation on the *CTCF* promoter is able to mask HREs to inhibit HIF1α binding, we performed HIF1α ChIP-qPCR in sgcontrol and dCas9-DNMT3A-sg2 cells under normoxic and hypoxic conditions. HIF1α-ChIP-qPCR showed that dCas9-DNMT3A-sg2 mediated HREs methylation abolishes HIF1α binding to the *CTCF* promoter (**Figure 4C**) and hence successful *CTCF* suppression of expression under hypoxia. These results were also nicely mirrored by likewise reduced levels of mesenchymal marker vimentin and increased expression of E-cad in dCas9-DNMT3A-sg2 cells in comparison to the sgcontrol under hypoxic conditions (**Figure 4D**). We next performed the invasion assay and found that dCas9-DNMT3A-sg2 cells exhibited reduced invasive phenotype as compared to sgcontrol cells under hypoxia corroborating with the reduced *CTCF* expression and its effects on the invasive phenotype of cells under hypoxia (**Figure 4E and 4F**). Ultimately, we found that *CTCF* expression can be modulated by epigenetic therapy using the CRISPR dCas9 system to render hypoxic cancer cells EMT phenotype.

**Figure 4:**
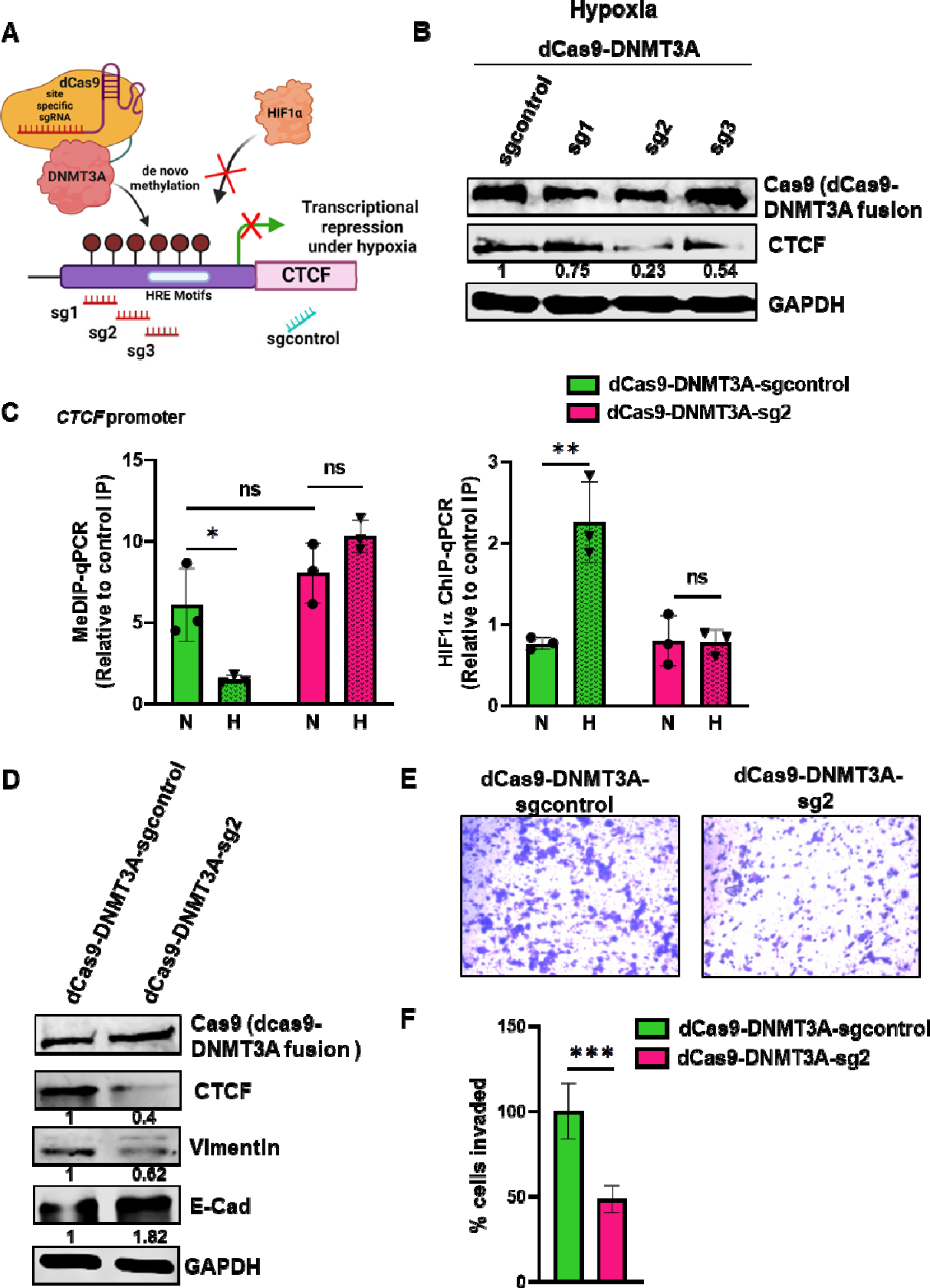
Epigenetic modulator dCas9-DNMT3A mediated HREs methylation is sufficient to suppress hypoxia-induced CTCF expression and impairs EMT phenotype. **A)** Schematic representation of targeting the CTCF promoter region by dCas9-DNMT3A with specific sgRNAs against HREs to maintain methylation and suppress CTCF expression **B)** Immunoblot of CTCF in MCF7 cells transfected with dcas9-DNMT3A-sgRNAs versus sgcontrol under hypoxia, GAPDH as loading control **C)** MeDIP-qPCR and HIF1α ChIP-qPCR in normoxic and hypoxic MCF7 cells transfected with dCas9-DNMT3A-sg2 versus sgcontrol **D**) Immunoblot of Cas9 (dCas9-DNMT3A fusion construct), CTCF, vimentin and E-cad in MCF7 cells transfected with dcas9-DNMT3A-sg2 in comparison to sgcontrol under hypoxia **E)** Invasion assay with **F)** quantification as % of cells invaded after MCF7 cells were transfected with either sgcontrol or dCas9-DNMT3A-sg2 under hypoxic condition. Error bar shows mean values ± SD. (n=3). As calculated using two-tailed Student’s t test, *p < 0.05, **p < 0.01, ***p < 0.001.

### *COL5A1* as a novel target of CTCF

It is well reported that hypoxia is a potent micro-environmental factor that promotes EMT and tumor metastasis ^2, 27^ and we have shown *CTCF* upregulation under hypoxia. These observations prompted us to hypothesize that CTCF might regulate the expression of EMT-related genes under hypoxia to render a metastatic phenotype. To decipher novel breast cancer-specific hypoxia-induced EMT gene regulated by CTCF, we first retrieved the list of genes upregulated in the breast tumor samples from GEPIA (**Table S1**) and then compared the list of gene upregulated in breast cancer with the hallmark EMT gene dataset to extract overlapping genes. Further, these genes overlapped with hypoxia hallmark geneset (MSigDB) to extract hypoxia-regulated breast cancer specific EMT genes (**Figure S5A**). Our ChIP-seq data revealed enriched CTCF binding sites in *COL5A1* promoter region under hypoxia (**Figure S5B**). The involvement of collagens in cancer progression has been reported recently ^30–32^. Despite their critical importance, only a few studies describe their functional role in cancer metastasis and their role in hypoxia driven cancer progression. Fibril-forming collagen, *COL5A1* ^33, 34^ is associated with pathological diseases such as classic Ehlers–Danlos syndrome and cancer ^35^. Hence, *COL5A1* being novel target of CTCF and was not previously studied in cancer hypoxic condition, was further selected as a model gene to study the transcriptional reprogramming in hypoxia-induced EMT genes that mediates invasiveness driven by CTCF.

### Hypoxia induced COL5A1 expression is both HIF1**α** and CTCF dependent in breast cancer cells

To validate the involvement of *COL5A1* in cancer progression, we first analyzed the *COL5A1* mRNA expression between tumor and normal tissues using the publicly available TCGA datasets available at GEPIA platform and found that the expression of *COL5A1* was up-regulated significantly in the tumor sample compared with the normal group in many cancers including breast cancer **(Figure 5A and Figure S6A)**. To examine whether *COL5A1* is induced in hypoxic tumor, we analyzed *COL5A1* mRNA expression in 165 tumor samples from a breast cancer patient microarray profile (GSE76250) ^36^ stratified as hypoxic_high or hypoxic_low based on the HYPOXIA_HALLMARK ^37^ gene signature using a method described previously ^6^. It appeared that the *COL5A1* is significantly upregulated (*p* = 3.08xe10^−8^) in samples stratified as hypoxic_high as compared to those stratified as hypoxic_low **(Figure 5B)**. Similar results were found when we analyzed the normoxic vs hypoxic HCC1806 HTA 2.0 array data from GEO (GSE147516) and observed significant upregulation of *COL5A1* in HCC1806 hypoxic cells **(Figure 5C)**. The results from these three independent cohorts indicated that *COL5A1* might possess a significant role in breast cancer progression under hypoxic condition.

**Figure 5:**
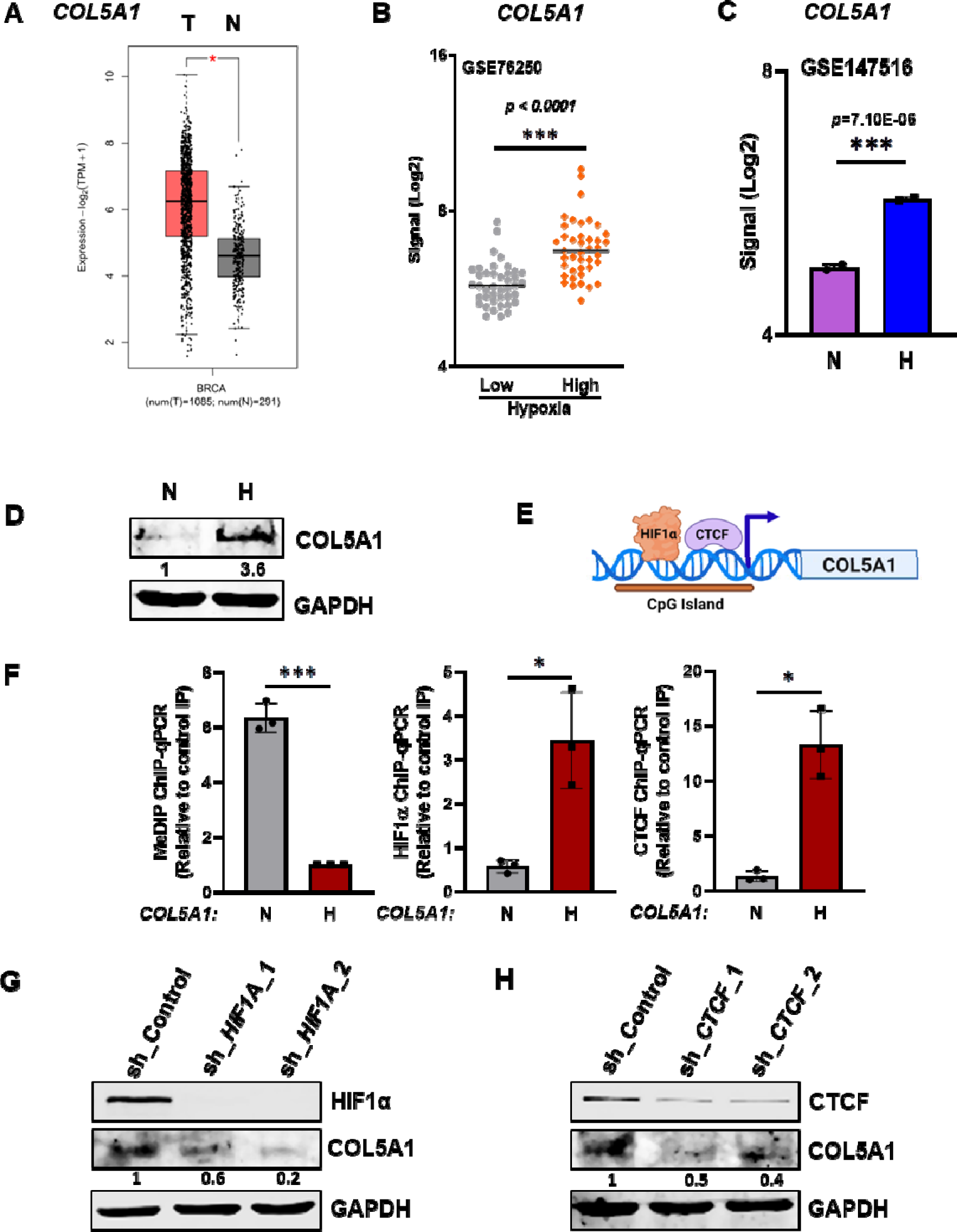
COL5A1 expression is induced under hypoxia in an HIF1α-CTCF-dependent manner. **A)** TCGA gene expression profile of COL5A1 pertaining to normal breast tissue (N) and primary breast tumor (T) **B)** COL5A1 mRNA expression for GSE76250 breast cancer dataset stratified as Hypoxic_high (n=41) or Hypoxic_low (n=41) **C)** Expression of COL5A1 in Microarray (using Human Transcriptome Array 2.0) data in HCC1806 (GSE147516) post hypoxic treatment **D)** Immno-blot of COL5A1 in normoxic and hypoxic breast cancer cell line, MCF7 **E)** schematic showing putative presence of HIF1α and CTCF on COL5A1 promoter **F)** MeDIP, HIF1α and CTCF ChIP-qPCR on COL5A1 promoter in normoxic and hypoxic MCF7 cells **G)** Immunoblot of COL5A1 in HIF1A knockdown **H)** Immunoblot of COL5A1 in CTCF knockdown and shControl MCF7 cells under hypoxia. Error bar shows mean values ± SD. (n=3). As calculated using two-tailed Student’s t test, *p < 0.05, **p < 0.01, ***p < 0.001.

To further strengthen these findings, we examined COL5A1 protein levels in breast cancer cell lines under normoxic and hypoxic condition. Immunoblot assay showed that COL5A1 protein levels were increased in both MCF7 and HCC1806, under hypoxic conditions (**Figure 5D and Figure S6B**). Taken together, these data indicate that hypoxia induces COL5A1 protein expression in human breast cancer. To further decode the hypoxia-mediated molecular mechanism regulating *COL5A1* expression under hypoxia, we examined *COL5A1* promoter and found putative HIF1α binding HRE motif and CTCF binding site in the promoter of *COL5A1* further suggests the involvement of these proteins in regulation of *COL5A1* expression (**Figure 5E and Table S2**). Promoter analysis also revealed the presence of dense CpG Island spanning HIF1α and CTCF binding sites and it is reported that binding of CTCF and HIF1α to genomic loci is inhibited by DNA methylation (**Figure 5E**) ^19, 21, 26^. Further, MeDIP-qPCR analysis revealed highly methylated DNA around TSS of *COL5A1* in normoxic cancer cells however, we observed reduced methylation levels on *COL5A1* promoter under hypoxic condition as compared to normoxic control (**Figure 5F and Figure S6C**). Furthermore, ChIP-qPCR of HIF1α and CTCF revealed enrichment of both HIF1α and CTCF on *COL5A1* promoter under hypoxia compared to normoxia (**Figure 5F and Figure S6C**). To determine whether HIF1α, CTCF, or both are required for hypoxia-mediated *COL5A1* expression, we silenced *HIF1A* or *CTCF* using shRNA in breast cancer cell lines (**Figure 5G and H and Figure S6D and S6E**).

Depleting HIF1α or CTCF, in breast cancer cells greatly diminished hypoxia-induced *COL5A1* expression (**Figure 5G and H and Figure S6D and S6E**). Together, these findings demonstrate that hypoxia-driven *COL5A1* expression is mediated by a tripartite mechanism involving epigenetic regulation, HIF1α and CTCF, collectively.

### Hypoxia drives alternative splicing event in *COL5A1* and favors exon 64A inclusion in breast cancer cells

Previous studies have identified an mutually exclusive alternative splicing event of exon 64 of *COL5A1* gene that give rise to isoform that either contain exon 64A or exon 64B ^38, 40^ (**Figure S6F**). Furthermore, mounting evidence suggests that one of the ways that cells adapt to hypoxia is through reprogramming the alternative splicing events ^4^. Therefore, we further expanded our study to delineate the hypoxia-driven COL5A1 exon 64 alternative splicing and its subsequent effect on breast cancer metastasis. To validate that hypoxia-drives alternative splicing of exon 64, we first analyzed the alternative splicing event in exon 64 in 165 tumor samples from a breast cancer patient microarray profile (GSE76250) stratified as hypoxic_low or hypoxic_high as described above. It appeared that the exon 64A is significantly included (p = 1.86 × 10−8) in samples stratified as hypoxic_high as compared to those stratified as hypoxic_low (**Figure 6A**). Similarly, HCC1806 HTA 2.0 array data from GEO (GSE147516) revealed significant inclusion (*p = 0.00002*) of exon 64A under hypoxia **(Figure 6B)**. The qPCR using exon-specific junction primers also showed a significant upregulation of exon 64A and downregulation of exon 6B isoform under hypoxia in breast cancer cell lines **(Figure 6C and Figure S6G)**. These findings indicated hypoxia-mediated alternative splicing of exon 64, and breast cancer cells specifically prefer the inclusion of exon 64A over exon 64B under hypoxia. Therefore, we sought to further define the molecular mechanisms driving *COL5A1* exon 64 alternative splicing under hypoxia.

**Figure 6:**
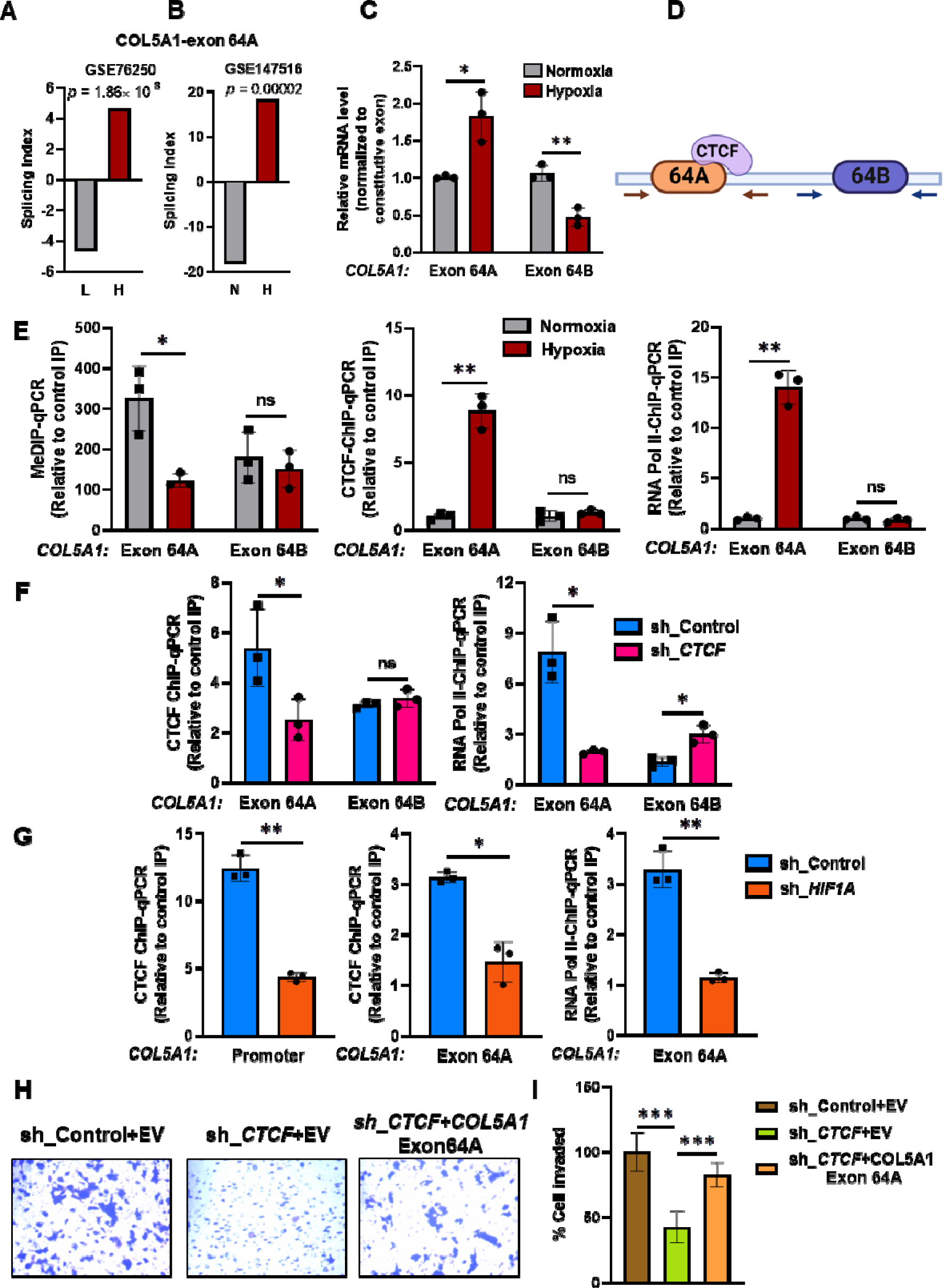
Hypoxia-induced COL5A1 exon 64AA inclusion is regulated by methylation dependent enrichment of CTCF on exon 64A and is associated with RNA Pol II pause. **A)** HCC1806 cells microarray data (GSE147516) showing significant inclusion of exon 64A of COL5A1 gene under hypoxia as compared to normoxia and is represented by splicing index values, **B)** significant inclusion of exon 64A of COL5A1 in GSE76250 breast cancer dataset stratified as Hypoxic_high (H, n=41) or Hypoxic_low (L, n=41) **C)** qRT-PCR analysis of COL5A1 exon 64A and exon 64B isoform normalized to RPS16 and constitutive exon expression levels in normoxic and hypoxic MCF7 cells **D)** Putative CTCF binding sites on exon 64A **E)**MeDIP, CTCF and RNA Pol II ChIP-qPCR for exon 64A and exon 64B in normoxic and hypoxic MCF7 cells **F)** CTCF and RNA Pol II ChIP-qPCR on exon 64A and exon 64B in MCF7 cells transduced with either shCTCF or shControl under hypoxia **G)** CTCF or RNA Pol II ChIP-qPCR on COL5A1 promoter or exon 64A in MCF7 cells transduced with either shHIF1A or shcontrol under hypoxia **H)** Invasion assay with **I)** its quantification as % of cells invaded after over expression of either empty vector or COL5A1exon 64A isoform in CTCF depleted MCF7 cells in comparison to the shControl cells under hypoxia. Error bar shows mean values ± SD. (n=3). As calculated using two-tailed Student’s t test, *p < 0.05, **p < 0.01, ***p < 0.001.

### Hypoxia-driven CTCF regulates the inclusion of exon 64A in *COL5A1* mRNA and promotes EMT in breast cancer cells

It is noteworthy to mention that CTCF is important for exon inclusion by pausing RNA Pol II-mediated transcription in a methylation-dependent manner ^19–21^. CTCF motif search revealed presence of putative CTCF binding motif immediately downstream to exon 64A **(Figure 6D and Table S2)**, corroborating with the ‘roadblock’ model where CTCF-mediated RNAPII elongation stalling occurs downstream of included exons ^21^. Therefore, we further investigated the role of CTCF in regulating hypoxia-mediated alternative splicing of *COL5A1* and the inclusion of exon 64A. First, we check the DNA methylation status of exons 64A and exon 64B using MeDIP-qPCR in normoxic vs hypoxic conditions. MeDIP-qPCR analysis revealed the presence of DNA methylation under normoxic condition on both exons, however, only exon 64A showed hypoxia-dependent reduced DNA methylation **(Figure 6E and Figure S7A)**. Further, CTCF-ChIP-qPCR analysis revealed CTCF enrichment on exon 64A and not on exon 64B in hypoxic condition as compared to the normoxic control **(Figure 6E and Figure S7A)**. We then validated RNA Pol II pause and performed RNA Pol II ChIP-qPCR. We found significant enrichment of RNA Pol II on exon 64A under hypoxic condition and no enrichment on exon 64B **(Figure 6E and Figure S7A)**. To further confirm the involvement of CTCF in mediating RNA Pol II pause, we performed RNA Pol II ChIP-qPCR in *CTCF* knockdown breast cancer cell lines under hypoxic condition. CTCF-ChIP-qPCR in CTCF-depleted breast cancer cell lines showed reduced enrichment of CTCF on exon 64A as compared to control cells under hypoxia **(Figure 6F and Figure S7B)**. Depletion of CTCF also reduced enrichment of RNA Pol II on exon 64A under hypoxia in compared to its hypoxic control cells **(Figure 6F and Figure S7B)**. Surprisingly, we observe a comparable shift in RNA Pol II enrichment on exon 64B in CTCF-depleted cells under hypoxia as compared to control hypoxic cells **(Figure 6F and Figure S7B)**. To validate that CTCF enrichment on exon 64A is due to hypoxia-driven CTCF expression regulated by HIF1α, we examined CTCF and RNA Pol II enrichment on exon 64A in HIF1α depleted cells under hypoxia. CTCF-ChIP-qPCR in HIF1α depleted cells showed reduced CTCF enrichment on *COL5A1* promoter **(Figure 6G)** and on exon 64A as compared to hypoxic control cells **(Figure 6G)**. RNA Pol II-ChIP-qPCR also showed reduced RNA Pol II enrichment on exon 64A in HIF1α depleted cells as compared to control cells under hypoxia **(Figure 6G)**. Collectively, hypoxia-mediated concomitant enrichment of CTCF on exon 64A and RNA Pol II pause let us state that *COL5A1* exon 64A inclusion is regulated by hypoxia-driven DNA de-methylation dependent CTCF-mediated RNA Pol II pause.

We further investigated the connection between hypoxia-driven CTCF-mediated inclusion of exon 64A and EMT phenotype of hypoxic breast cancer cells and performed invasion assay in CTCF depleted cells either over-expressed with *COL5A1*exon64A isoform or empty vector backbone under hypoxia. The rescue experiment showed that restoration of *COL5A1*exon 64A expression in CTCF-depleted breast cancer cell lines partially reversed the invasive-inhibiting effects of CTCF knockdown in comparison to the control cells under hypoxia (**Figure 6H and 6I and Fig S7C and S7D**). Accordingly, this demonstrated that hypoxia-induced CTCF expression modulates gene expression and alternative splicing event to drive the EMT phenotype of breast cancer cells.

### Targeted de Novo methylation of CTCF binding site at exon 64A alters CTCF-mediated *COL5A1* exon 64 alternative splicing

The previous experiments established a correlation between hypoxia-driven DNA-de-methylation and CTCF-mediated exon 64A inclusion in *COL5A1* mRNA. To test whether DNA methylation and CTCF enrichment were sufficient to drive alternative splicing of *COL5A1* exon 64, it was necessary to experimentally manipulate CTCF occupancy at exon 64A alone. To accomplish this, we used nuclease-deficient Cas9 (dCas9)-DNA methyltransferase, DNMT3A epigenetic editing system, for targeted DNA methylation of CTCF binding on exon 64A of *COL5A1* (**Figure 7A and Figure S7E**). We designed exon 64A-specific gRNAs targeting various sites across exon 64A, including CTCF site **(Figure 7A)**. To measure dCas9-DNMT3A-mediated-DNA methylation-driven alternative splicing switch, we first performed qRT-PCR using exon-specific primers in sgcontrol vs sgRNA specific to exon 64A under hypoxia.

**Figure 7:**
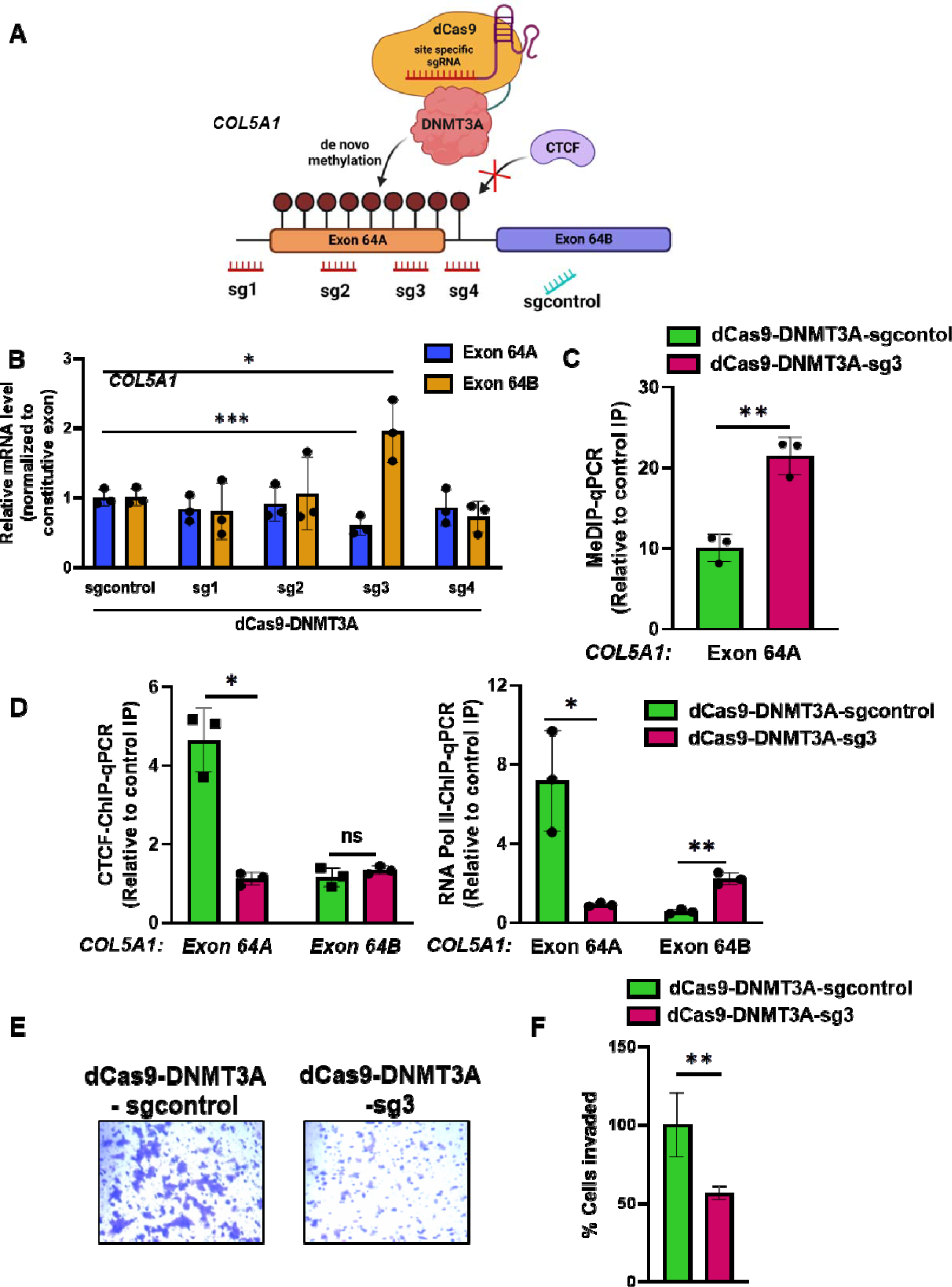
Targeted methylation of CTCF binding site on exon 64A by dCas9-DNMT3A block CTCF enrichment and reduced inclusion of exon 64A in COL5A1 mRNA under hypoxia. **>A)** Schematic representation of targeting the putative CTCF binding sites on COL5A1 exon 64A by dCas9-DNMT3A with specific sgRNAs to maintain methylation under hypoxia to suppress CTCF enrichment **B)** qRT-PCR of COL5A1 exon 64A and exon 64B in MCF7 cells transfected with dCas9-DNMT3A-sgRNAs or sgcontrol under hypoxia **C)** MeDIP-qPCR on exon 64A **D)** CTCF ChIP-qPCR and RNA Pol II ChIP-qPCR on exon 64A and exon 64B in hypoxic MCF7 cells transfected with dcas9-DNMT3A-sg3 versus sgcontrol **E)** Invasion assay with **F)** quantification as % of cells invaded after MCF7 cells were transfected with either sgcontrol or dcas9-DNMT3A-sg3 under hypoxic condition. Error bar shows mean values ± SD. (n=3). As calculated using two-tailed Student’s t test, *p < 0.05, **p < 0.01, ***p < 0.001.

The dCas9-DNMT3A mediated epigenetic editing significantly switch the splicing events of exon 64 in dCas9-DNMT3A-sg3 (situated upstream to CTCF binding site immediately downstream to the exon 64A) and not in other sgRNAs targeting other exon 64A sites as compared to the sgcontrol under hypoxia **(Figure 7B)**. The mRNA levels of exon 64A reduced significantly while of exon 64B is upregulated in dCas9-DNMT3A-sg3 vs sgcontrol under hypoxia **(Figure 7B)**.

To confirm that the decreased expression of exon 64A is due de novo methylation and blocking of CTCF enrichment, we performed MeDIP-qPCR and CTCF ChIP-qPCR in cells transfected with dCas9-DNMT3A-sg3 and sgcontrol under hypoxia. MeDIP-qPCR showed that dCas9-DNMT3A-sg3 increased significant methylation on exon 64A compared to the sgcontrol (**Figures 7C**). Further, CTCF-ChIP-qPCR in dCas9-DNMT3A-sg3 transfected cells showed significant reduction in CTCF enrichment on exon 64A as compared to the sgcontrol under hypoxia (**Figures 7D**). Similarly, RNA Pol II-ChIP-qPCR showed significantly reduced RNA Pol II enrichment on exon 64A and increased enrichment on exon 64B in dCas9-DNMT3A-sgRNA3 as compared to the sgcontrol under hypoxia (**Figures 7D**), supporting the notion that DNA methylation blocks CTCF binding and thus alters the *COL5A1* exon 64 alternative splicing outcomes in hypoxic condition. A similar set of experiment was performed to assess the effect on invasive phenotype of the cells treated either with dCas9-DNMT3A-sgRNA3 or sgcontrol and result presented in **Figure 7E and 7F** demonstrated the decreased in cell invasion number in dCas9-DNMT3A-sgRNA3 in comparison to the sgcontrol. These data revealed that DNA methylation regulated enrichment of CTCF and RNA Pol II pause at exon 64A under hypoxia is sufficient to drive AS of exon 64 of *COL5A1* and that reduced exon 64A inclusion greatly impaired EMT phenotype of the breast cancer cell under hypoxia.

### CTCF-mediated promoter-exon upstream looping regulate CTCF-mediated Pol II pause at exon and inclusion of *COL5A1* exon 64A under hypoxia

One of the alluring observations made during this study is that COL5A1 gene is 205 kb long with a mutually alternative splicing event present for last 3^rd^ exon situated 188 kb downstream of the promoter. Besides, CTCF-ChIP-qPCR showed direct binding of CTCF on *COL5A1* promoter as well as exon 64A under hypoxia. Given the evidence that CTCF-mediated promoter-exon loop formation drive the alternative splicing of exons situated far from the promoter ^39^, we next asked a question whether CTCF binding at the promoter and proximal to exon 64A plays any role in bringing exon 64 into close physical proximity with its promoter and regulating exon 64 alternative splicing under hypoxia. It is well known that CTCF-mediated chromatin loops preferentially form between two convergently bound CTCF molecules and sense CTCF motif bias at promoter or anchor site is also evident ^39, 41, 42^. The CTCF motif at *COL5A1* promoter that showed enrichment under hypoxia **(Figure 5F)** is in sense orientation corroborating with the previous studies. We then scanned the upstream and downstream proximity of exon 64 to find antisense CTCF motif to assess the potential intragenic CTCF-mediated promoter-exon looping. We found 3 CTCF motif (p<0.01 or 0.0001) in antisense orientation in the proximity of exon 64 **(Table S2)**.

To investigate whether there is a long-range DNA-looping interaction between *COL5A1* promoter and CTCF motifs proximal to exon 64 we performed chromosome conformation capture (3C) assays under normoxia and hypoxia condition. We used primer against promoter as anchor primer and CTCF motifs proximal to exon 64 as test primer **(Figure 8A)**. When using promoter anchor primer, strong interaction was detected with the test primer T1 corresponding to CTCF motif situated ∼7kb upstream from exon 64A under hypoxia when compared to the normoxic cells whereas with the test primer T2 (CTCF motif situated ∼5.6kb upstream from exon 64A) and T3 (CTCF motif situated ∼700bp downstream from exon 64A) interactions were not detected (**Figure 8B**). Further, in CTCF KD hypoxic breast cancer cell, we observe less interaction between promoter anchor primer and Test primer T1 in 3C analysis (**Figure 8B**). To confirm the involvement of CTCF in bringing promoter-upstream exon loop, CTCF ChIP-qPCR experiment was done and showed CTCF enrichment at CTCF motif situated ∼7kb upstream of exon 64A under hypoxia corresponding to the test primer T1 whereas no enrichment were found at CTCF motif corresponding to T2 and T3 when compared to normoxic cells **(Figure 8C)** confirming CTCF-mediated promoter-upstream exon looping under hypoxia.

**Figure 8:**
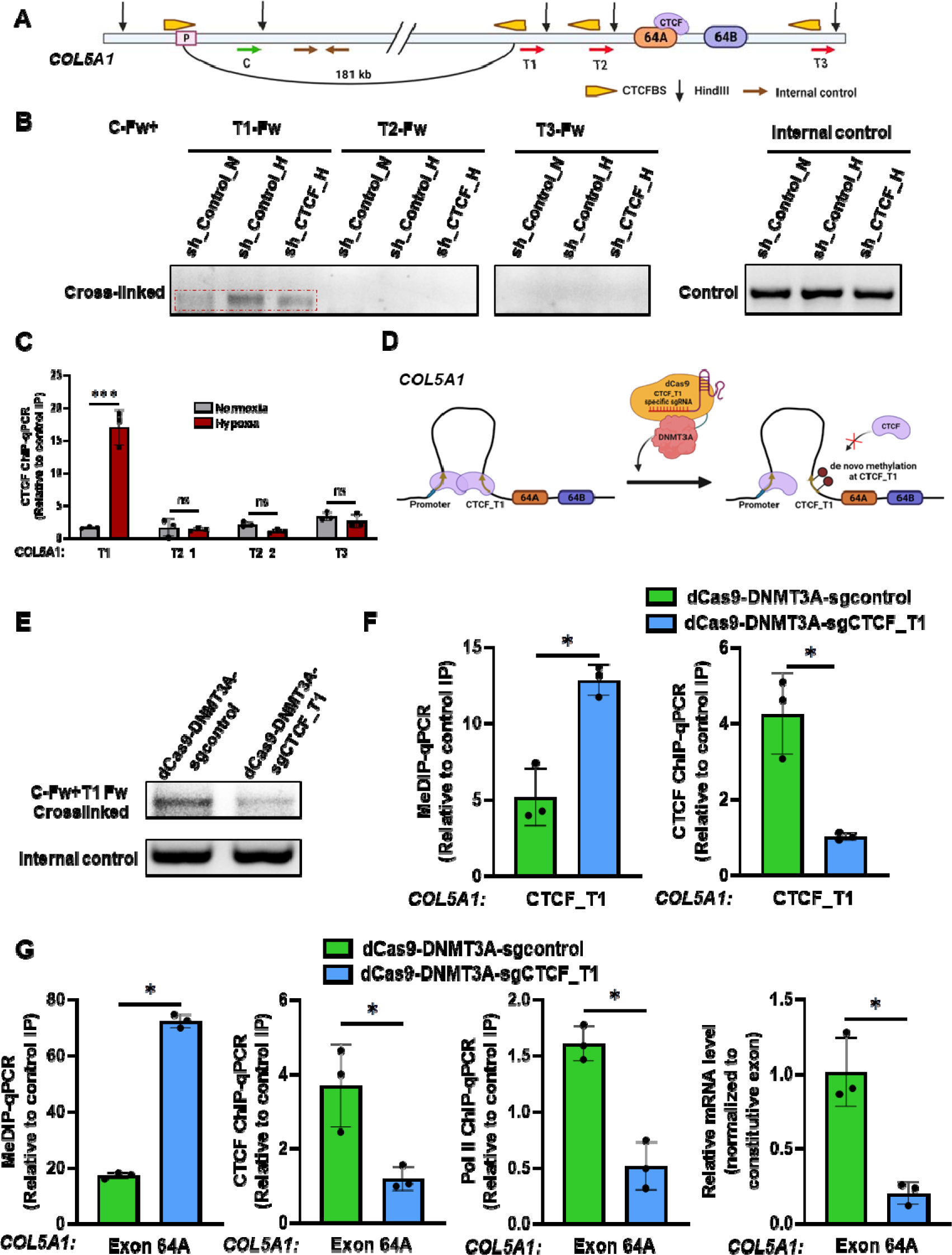
Hypoxia-induced CTCF mediated COL5A1 promoter-exon upstream looping decides inclusion of COL5A1 exon 64A by regulating CTCF-mediated Pol II pause at exon 64 under hypoxia. **A)** Schematic depicts genomic view of COL5A1 gene loci on chromosome 9 and various putative convergent CTCF sites present in the promoter and proximal to exon 64. The positions of the CTCF recognition (yellow), HindIII sites (black arrow), and primers are indicated. C indicates the 3C anchor site and T1, T2 and T3 indicate test sites **B)** 3C analysis showing 3C assay for DNA-looping interactions between different HindIII fragments using an anchor primer C with three test primers T1-T3 showing formation of COL5A1 promoter-exon upstream 181 kb long loop mediated by T1 test site upstream of alternatively spliced exon 64 **C)** CTCF enrichment was measured by ChIP-qPCR at putative convergent CTCF motif (T1-T3) proximal to exon 64 in normoxic and hypoxic MCF 7 cells **D)** Schematic representation of targeting the CTCF binding site involved in looping by dCas9-DNMT3A with specific sgRNAs to induce de novo methylation to open CTCF loop **E)** 3C analysis of cells transfected with dcas9-DNMT3A-CTCF_T1 or sgcontrol using anchor primer C and test primer T1 under hypoxia **F)** MeDIP and CTCF ChIP-qPCR in cells transfected with dcas9-DNMT3A-sgCTCF_T1 versus sgcontrol hypoxic MCF7 **G)** MeDIP, CTCF and RNA Pol II ChIP-qPCR on exon 64A and exon 64A qPCR to check its inclusion in MCF7 cells transfected with dcas9-DNMT3A-CTCF_T1 versus sgcontrol under hypoxia. Error bar shows mean values ± SD. (n=3). As calculated using two-tailed Student’s t test, *p < 0.05, **p < 0.01, ***p < 0.001.

Taken together, these results reveal that CTCF participates in the long-range chromatin interaction that is ∼181 kb long and brings together the COL5A1 promoter and exon 64A in close proximity via intragenic CTCF **(Figure 8A)**.

To define the correlation between alternative splicing events and CTCF-mediated promoter-intragenic looping hypoxia, we employed dCas9-DNMT3A mediated DNA methylation at intragenic CTCF binding site (corresponding to T1) to restrict CTCF binding and disruption of promoter-upstream exon loop. We designed specific sgRNAs **(Figure 8D)** targeting dCas9-DNMT3A to CTCF_T1 site to investigate whether de novo methylation would interfere with the looping function of CTCF. 3C analysis showed a significant reduction in interaction between promoter anchor primer and Test primer T1 in dCas9_DNMT3A-CTCF_T1 comparison to the sgcontrol under hypoxia **(Figure 8E)**. MeDIP-qPCR analysis revealed increased DNA methylation at T1-CTCF binding site and corroborates with decreased CTCF enrichment under hypoxia at T1-CTCFBS as analyzed by CTCF-ChIP-qPCR in dCas9_DNMT3A-CTCF_T1 in comparison to the control **(Figure 8F)**. As stated above, inclusion of exon 64A is regulated by hypoxia-driven DNA de-methylation dependent CTCF-mediated RNA Pol II pause we next sought to answer how looping disruption affects all these events for exon 64A under hypoxia. To answer this, we investigated DNA methylation status, CTCF and RNA Pol II enrichment on exon 64A in dCas9_DNMT3A-CTCF_T1 in comparison to the sgcontrol. MeDIP-qPCR showed increased DNA methylation on exon 64A, moreover, CTCF and RNA Pol II ChIP-qPCR showed reduced enrichment on exon 64A in dCas9_DNMT3A-CTCF_T1 in comparison to the sgcontrol under hypoxia **(Figure 8G)**. qPCR analysis showed significant reduction in the inclusion of exon 64A under hypoxia in dCas9_DNMT3A-CTCF_T1 in comparison to the sgcontrol **(Figure 8G)**. Collectively, we proposed a model where hypoxia-driven *COL5A1* AS is regulated by an intricate mechanism coupling epigenetics with CTCF-mediated promoter-exon upstream looping and RNA Pol II pausing at exon 64A.

### Hypoxia-driven CTCF differential binding regulates transcriptional and alternative splicing reprogramming in hypoxic breast cancer cells

To investigate the impending mechanisms of CTCF regulation of transcription and alternative splicing in hypoxic condition, we performed CTCF ChIP-seq in normoxic and hypoxic MCF7 cells. We identified 24104 CTCF sites were significantly gained from normoxia to hypoxia (**Figure 9A**). We identify 3040 genes overlapped with the signals of gained CTCF peaks within ±5_kb elements proximal to transcriptional start sites that might be directly targeted by hypoxia-driven differential CTCF occupancy. Gene ontology analysis revealed that CTCF-dependent transcripts were strongly enriched for processes related to cell proliferation, EMT, stemness and cancer-related pathways (**Figure 9B**). To determine the effect of CTCF gained occupancy on hypoxic cell transcriptome diversity, we intersected the genes upregulated in normoxic vs hypoxic condition (GSE166203) obtained using NOIseq with CTCF peaks significantly gained in hypoxia and identified 408 genes (**Figure 9C**) protein-coding showing enrichment of genes involved in cell cycle, EMT, hypoxia, and cancer pathways (**Figure 9D**).

**Figure 9:**
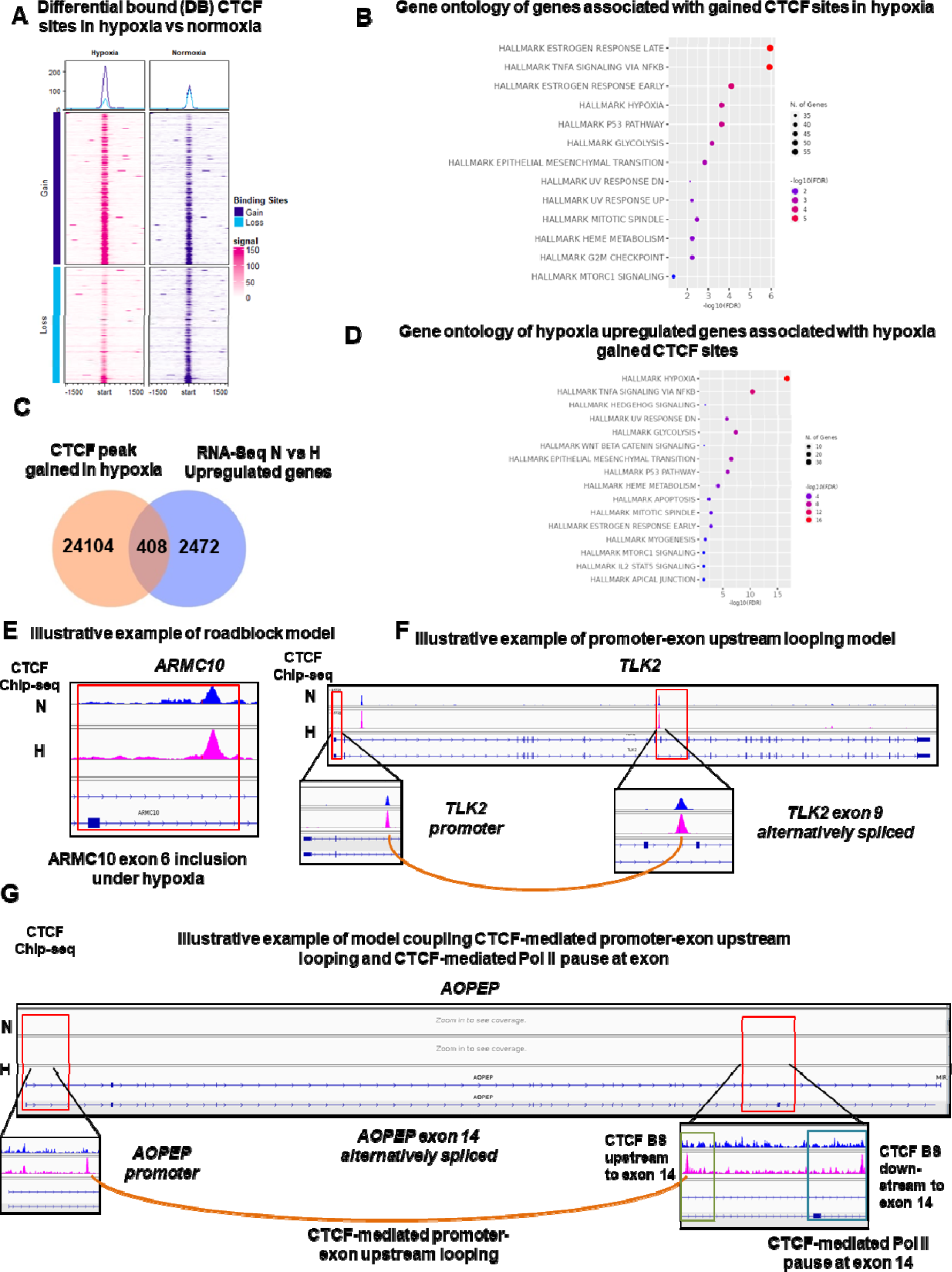
Hypoxia-induced differential CTCF occupancy regulates transcriptional and alternative splicing dynamics in breast cancer cells to drive EMT. **A)** Differential binding analysis identified 24104 CTCF binding sites significantly gained under hypoxia in MCF7 cells (p <0.05). Top 1000 differential bound sites are shown **B)** Gained CTCF binding sites were significantly enriched within the promoters of many genes and representative significantly enriched terms from the Gene Ontology and curated MsigDB **C)** Venn diagrams showing the overlap between hypoxia vs normoxia gained CTCF sites (from **A**) and hypoxia vs normoxia upregulated genes (from GSE166203). **D)** Representative significantly enriched terms from the Gene Ontology and curated MsigDB highlights EMT related pathways **E)** illustrative examples of exon inclusion under hypoxia by roadblock model **F)** illustrative examples of exon inclusion under hypoxia by promoter-exon upstream looping model **G)** illustrative examples of exon inclusion under hypoxia by model coupling CTCF-mediated promoter-exon upstream looping and CTCF-mediated Pol II pause at exon showed in the present study.

Showing the importance of CTCF in shaping alternative splicing programs, particularly exon inclusion, we next investigated the relationship between significantly gained CTCF site and alternative splicing events occurring in hypoxic cells. For this, we first analyzed the alternative splicing events in normoxic vs hypoxic conditions (GSE166203) and found that 18826 many exon inclusion events present in hypoxic condition (**Figure S8A**). To find CTCF mediated exon inclusion events under hypoxia, we quantified the distribution of hypoxia-gained CTCF sites with exons included under hypoxia and found that 2017 events overlapped with hypoxia-gained CTCF binding sites and possibly regulate exon inclusion under hypoxia.

We also found that genes such as *CUX1, RNF150 and ITPR1* are perhaps regulated by model proposed in the present study coupling CTCF-mediated promoter-exon upstream looing and CTCF-mediated RNA Pol II pause at differentially used exon. The examples illustrate of each group is shown in **Figure 9E, 9F and 9G** and **Figure S9B, S9C and S9D**. A gene ontology analysis on the genes containing CTCF-regulated exons showed a strong enrichment for genes related to cell motility, EMT pathways and cancer metastasis. Overall, this analysis strengthens the model presented in this study integrating the multiple layer of alternative splicing regulated by CTCF that promotes EMT in breast cancer cells under hypoxic condition.

## Discussion

Rapidly growing solid tumor outstrips the development of new blood vessels and creates a hypoxic niche in the core, which is a critical factor for cancer progression. Hypoxia elicits epithelial−mesenchymal transition (EMT) to drive the initiation of metastasis by modulating the expression and alternative splicing of a large array of genes as well as by regulating the epigenetic landscape of the genome ^3, 4, 7, 11, 27, 28, 43^. However, within each cancer, the specific target genes activated and the pathogenic consequences of their activation are unique. Here, we report that hypoxia, through epigenetic modulation, induces *CTCF* expression in breast cancer. We have demonstrated the critical role of epigenetic marks specifically DNA methylation in regulating *CTCF* expression in breast cancer. The *CTCF* promoter contains dense CpG Island which is highly methylated under normoxic condition. DNA methyltransferase, DNMT3A is responsible for maintaining the methylation of CTCF promoter under normoxic condition. Indeed, we found a positive role of DNA methylation in driving CTCF expression as depletion of DNMT3A reduces DNA methylation on *CTCF* promoter, which led to the reduced *CTCF* expression in breast cancer cells in normoxic condition. Previously, DNA methylation in very few studies have been reported as a positive regulator of gene expression ^24^ and decoding the molecular switch regulating *CTCF* expression through DNA methylation will be an important area of further study.

In the present study, we have shown that *CTCF* is a direct target of HIF1α. Hypoxia-induced DNA de-methylation at CpG Island gives access to the HRE motifs present in the *CTCF* promoter and thereby allows HIF1α recruitment. We observed hypoxia-associated increase in epigenetic marks H3K4me3 and H3K27ac on CTCF promoter around HRE element consistent with the previous study showing a selective increase of these marks by HIF complex to facilitate rapid activation of HIF target genes ^25^. Further, HIF1α depletion abrogates *CTCF* induction under hypoxia in breast cancer cell lines. We also uncover an intriguing process and showed that hypoxia-induced DNA de-methylation at *CTCF* promoter is HIF1α independent and HIF1α can only bind to un-methylated HRE elements. Our findings reinforce a growing body of evidence that highlights that DNA methylation repel HIF1α binding ^26^. We further showed hypoxia-induced EMT in breast cancer cell lines were drastically reduced by the *CTCF* depletion. Hypoxia-induced EMT was rescued in HIF1α-depleted cells by the restoration of CTCF expression, suggesting hypoxia-induced HIF1α-CTCF axis is one of the major drivers of EMT. When the HRE elements of *CTCF* promoter were targeted by dCas9-DNMT3A, extensive methylation of *CTCF* promoter was observed under hypoxia, which annulled HIF1α binding and hence suppression of hypoxia-induced CTCF expression. This targeted DNA methylation of *CTCF* promoter, in turn obliterated hypoxia-induced EMT. Taken together, our findings delineate a molecular mechanism by which *CTCF* expression is regulated under hypoxia through a critical signal transduction pathway that triggers firstly, DNA de methylation at *CTCF* promoter and secondly, HIF1α-mediated CTCF induction that promotes breast cancer cell EMT, a first step in metastasis. Ultimately, precisely controlled modulation of hypoxia-induced *CTCF* expression by dCas9-DNMT3A system depicts epigenetic therapeutic settings to render hypoxic cancer cells EMT phenotype.

Further, analysis showed genes that are upregulated in breast cancer and are involved in EMT and hypoxia, *COL5A1* was found to be a novel direct target of CTCF. Further TCGA data analysis showed increased expression of *COL5A1* in tumor samples as compared to the normal breast tissue. *COL5A1* also showed increased expression in tumor samples (GSE76250) stratified as hypoxia_high in comparison to the hypoxia_low. *COL5A1* also showed induced expression in the hypoxic condition in HCC1806 breast cancer cell line HTA 2.0 array data from GEO (GSE147516). We also showed that COL5A1 protein expression is induced under hypoxia in breast cancer cell lines, MCF7 and HCC1806. We found enrichment of both CTCF and HIF1α on *COL5A1* promoter under hypoxia and also observed reduced DNA methylation at *COL5A1* promoter consistent with the previous report showing a negative correlation between DNA methylation and CTCF/HIF1α binding ^21, 26^. We showed that both CTCF and HIF1α depletion abolished *COL5A1* expression under hypoxia, suggesting that these proteins might work together to regulate a distinct set of genes in the metastatic process under hypoxia. Although, further studies are required to more rigorously determine the role of HIF1α-CTCF complex in hypoxia-mediated metastasis.

Additionally, we also reported hypoxia-induced alternative splicing event in exon 64 for *COL5A1* transcript consistent with the previous studies that have identified alternative splicing of exon 64 of *COL5A1* gene ^38, 40^. We showed that hypoxic breast cancer cells prefer inclusion of exon 64A in *COL5A1* mRNA. An outstanding question in splicing biology concerns the mechanism of splicing specificity. Recent studies, including our own, showed CTCF as a regulator of alternative splicing ^9, 19–21, 39^. We also observed that *COL5A1* is a 205kb long with number of CTCF binding motifs across the gene. Indeed, we found CTCF binding motif immediately downstream to the exon 64A, suggesting CTCF as a possible key regulator of exon 64 alternative splicing. We observed reduced DNA methylation on exon 64A and not on exon 64B moreover, we found CTCF and RNA Pol II enrichment on exon 64A only in hypoxic cells corroborating with the previous studies that DNA methylation is one of the key factor deciding the binding of CTCF to its target loci ^19, 21^. Further experiments confirmed that CTCF enrichment on *COL5A1* promoter and exon 64A under hypoxia is HIF1α dependent. The dCas9-DNMT3A-mediated targeted DNA methylation of CTCF binding site present immediately downstream to exon 64A under hypoxia abolishes CTCF enrichment and sufficiently drive the switch in alternative splicing of exon 64 where it favors the inclusion of exon 64B instead of exon 64A. we also confirm that overexpression of COL5A1exon64A isoform restores the levels of invasiveness in the CTCF depleted hypoxic breast cancer cells, supporting a critical role of CTCF-COL5A1exon 64A axis in hypoxia-mediated tumor invasion. Here, we provide strong evidence that hypoxia-induced CTCF enrichment-mediated RNA Pol II pause favors exon 64A inclusion in COL5A1 mRNA that promotes EMT.

Till now, most studies have demonstrated the role of DNA methylation and CTCF-mediated RNA Pol II elongation related mechanisms. Recently, the involvement of intragenic CTCF binding sites, particularly those present proximal upstream to splice junctions in deciding alternative exon or intron inclusion in pre-mRNA is been elucidated ^19–21, 39, 44, 45^. Yet the interconnection between the two CTCF-mediated mechanisms regulating AS, particularly influence of chromatin architecture on CTCF-mediated RNA Pol II pause and exon inclusion remain largely uncharacterized. Here, we expanded our study to particularly decode the influence of chromatin architecture on CTCF-mediated RNA Pol II pause and exon inclusion as first, our model gene *COL5A1* is 205kb long with alternatively spliced exon is situated far from promoter (181kb), and second, we observed CTCF enrichment under hypoxia on *COL5A1* gene promoter as well as on proximal upstream to alternatively spliced exon 64, together suggests possible promoter-exon upstream-mediated long-range chromatin looping. Our data demonstrated the formation of promoter-exon upstream CTCF-mediated chromatin looping in *COL5A1* gene under hypoxia. Disruption of the this looping by targeted DNA methylation by dCas9-DNMT3A with specific sgRNA against intragenic CTCF binding site involved in looping inhibited DNA de methylation on exon 64A under hypoxia. This annulled CTCF enrichment on exon 64A and RNA Pol II pause, hence inclusion of exon 64A under hypoxia. Altogether, the results here demonstrate that intragenic CTCF-mediated promoter-exon upstream looping influence DNA methylation, CTCF enrichment and RNAPII pause at alternatively spliced exon and hence decide CTCF-regulated exon inclusion. However, while there is clear evidence that CTCF-mediated promoter-exon upstream looping affects CTCF-mediated RNA Pol II pause at alternatively spliced exon, identification of novel splicing mechanisms at the chromatin level is beyond the scope of this study, it will be of future interest to explore the CTCF-mediated promoter-exon chromatin looping associated splicing machinery.

The data presented in the present study provides new insights into hypoxia-CTCF axis mediated regulation of metastatic genes. It is the first report that shows that *CTCF* is induced under hypoxia through epigenetic regulation and is a target of HIF1α. Global analysis using CTCF ChIP-seq and RNA seq data showed that CTCF possibly regulates many gene expression and alternative splicing events related to metastatic genes. In continuation, we experimentally validated the role of CTCF in regulating alternative splicing of *COL5A1* under hypoxia. In conclusion, a hypothetical model is proposed that demonstrates the hypoxia-CTCF axis regulating the expression of EMT genes to ensure cancer cell survival and epithelial to mesenchymal transition that drives metastasis. Further investigations will provide a better understanding of CTCF regulated pathways that modulate cancer metastasis, such as epithelial to mesenchymal transition (EMT), adaptive metabolic reprogramming, angiogenesis and invasion. Owing the crucial role of cancer metastasis in patient prognosis, this study will aid in designing of new therapeutic strategies. At last, we employ epigenome editing strategies for potent, specific, and stable disruption of HIF1α or CTCF binding hence HIF1α-CTCF-COL5A1exon 64A axis that alleviates EMT potential of breast cancer cells under hypoxia and may represent a novel therapeutic target in breast cancer.

## Acknowledgments

P.K. is a recipient of Post-doctoral fellowship from Indian Institute of Science Education and Research Bhopal, India and Department of Biotechnology (DBT, India), Research Associate fellowship award. S.D. is a supported by Department of Biotechnology (DBT, India). D.P. is a recipient of funding from University Grants Commission, India. This work is supported by DBT/Wellcome Trust India Alliance Fellowship Grant IA/I/16/2/502719 (to S.S.).

## Author Contributions

S.S and P.K. designed the study; P.K. performed all experiments, generated data, and compiled figures; P.K. and S.D. performed luciferase assays and cloning; P.K. and D.P. analyzed ChIP-seq and RNA–seq data; P.K. and S.S. wrote the manuscript with inputs from all other authors.

## Data and Software Availability

CTCF ChIP-Seq data is deposited at GEO database and awaiting accession number. This study does not report any original code.

