## Supplementary information for "Hypoxia-induced CTCF mediates alternative splicing via coupling chromatin looping and RNA Pol II pause to promote EMT in breast cancer"

$ Present affiliation: Neuberg Supratech Kotgirwar Diagnostic, Bhopal.

**Material and Methods**

**Cell culture and treatment**

Human breast cancer cell lines MCF7 and HCC1806 were obtained from American Type Culture Collection (ATCC) and were cultured in ATCC recommended media DMEM for MCF7 and RPMI1640 for HCC1806 supplemented with 10% fetal bovine serum (FBS; Sigma,F7524), 100 units/ml of penicillin and streptomycin (Invitrogen, 15140122) and 2 mmol/lL-glutamine (Sigma, G7513). Cells were cultured at 37^0^C and 5% CO2. For treatment under hypoxic conditions (1% O2), cells were kept in a Ruskinn INVIVO2 400 hypoxia chamber.

**Breast cancer sample collection and Immunohistochemistry**

The study was approved by the Institute Ethics Committee of the Indian Institute of Science Education and Research Bhopal, India. Formalin-fixed, paraffin-embedded human breast cancer tissue sections were obtained from Bansal Hospital, Bhopal, India. Informed consent was obtained from all the patients. Clinical characteristics of patients used in the study are presented in **Supplementary Table S4**. Firstly, slides were fixed at 65°C for 2 hrs on the hot plate, deparaffinized and rehydrated as described previously ^1,2^. Subsequently, heat induced antigen retrieval was performed using 10 mM sodium citrate buffer (pH 6 in the laboratory microwave for 14 mins). Endogenous peroxidase was quenched with 1:10 dilution of 3% hydrogen peroxidase in methanol, followed by blocking with 3% bovine serum albumin (BSA). Primary antibodies against CAIX (1:50), and CTCF (1:200) were used (details of the antibodies are provided in **Supplementary Table S5**). Further, HRP/DAB-chromogenic based Super Sensitive™* Polymer-HRP Detection System kit (BioGenex, Catalog no. QD430-XAKE) was used as per manufactures’ instructions. All slides were counterstained with Harris’ hematoxylin (Merck). The images were captured by Thermo Scientific™ Invitrogen™ EVOS™ FL Auto 2 Imaging System and at 10× and 40× magnifications. Images were then processed in Adobe Photoshop Version 7.0. Quantification was done using color deconvolution in ImageJ (Fiji) and mean gray values were then converted to optical density.

**Generation of CTCF promoter construct**

To generate promoter construct, human CTCF (transcript, NM_006565) promoter sequence was retrieved from the Eukaryotic Promoter Database (<https://epd.vital-it.ch/>, ^3^). The CTCF gene promoter from −687 bp upstream to +145 bp downstream of the TSS at +1 was amplified by PCR using MCF7 genomic DNA as a template and inserted in the pGL3-Basic expression vector (Promega). Primers are given in **Supplementary Table S6**. CTCF promoter construct was cloned between the KpnI F and HindIII R sites.

**Site-directed mutagenesis**

The site-directed mutant construct of the CTCF promoter-luciferase construct was prepared using oligonucleotides with mutations in the HRE site present from +255 bp and +247 bp upstream of the CTCF TSS (GGCGTG to GTTTTG). The wild-type -687 to +145 CTCF promoter luciferase construct was used as a template. The SDM primers are given in **Supplementary Table S6**. The Mutations in the CTCF promoter was confirmed by DNA sequencing.

**Luciferase reporter assays**

MCF7 or HCC1806 cells (0.05 ×10^6^ or 0.3×10^6^) were seeded in 24-or 6-well plates, respectively and cultured for 16 h. The cells were co-transfected with CTCF promoter (-687 to +145) –luciferase construct and pRL-TK Renilla luciferase plasmid (Promega, E2231). After 12 h of transfection, hypoxic treatment was given and cells were lysed in passive lysis buffer (1% Triton-X-100, 25mM Tricine pH 7.8, 15mM potassium phosphate pH 7.8, 15mM magnesium sulphate, 4mM EGTA, 1mM DTT). The firefly luciferase activities were measured in a SpectraMax-Multi Detection System, and the values were normalized to Renilla luciferase activities. The relative values are represented as mean ± SD of triplicates.

**Molecular cloning of full length over expression constructs**

The CTCF and COL5A1 exon64A isoform CDS were cloned in pCMV-3Tag1A (Agilent, 240195) and pEGFP-C3 (Addgene, 6082-1) overexpression plasmids, respectively using MCF7 cDNA as a template and amplified using phusion DNA polymerase (NEB, M0530S). Primers are given in **Supplementary Table S6**. CTCF was cloned between the BamHI F and HindIII R sites and COL5A1 exon64A isoform was cloned between the HindIII F and KpnI R sites.

**Genomic DNA isolation and Methylated DNA immunoprecipitation (MeDIP-qPCR)**

After treatment, treated and control cells were processed to obtain gDNA using gDNA isolation kit (Sigma, G1N70) following manufacturer’s instructions. Methylated DNA immunoprecipitation (MeDIP) was performed as described before ^4,5^. Briefly, 5 μg of gDNA was sonicated to a fragment size of ~100–400 bp using Bioruptor sonicator (Diagenode), heated at 95^0^C for 10 mins and then incubated with 5-methyl cytosine antibody and normal rabbit IgG antibody (antibody details in **Supplementary Table S5**) for overnight at 4^0^C followed by isolation of Antibody/DNA complexes using Dynabeads Protein G (Invtrogen, Cat No. 10004D) hybridized with antibody. The obtained complex and input DNA were reverse cross-linked in TE buffer with 1% SDS and proteinase K (Invitrogen, 25530049, lot no. 2291039) for overnight at 65^0^C. Eluted DNA was then purified using PCR purification kit (QIAGEN, 28106). Immunoprecipitated fractions and 5% input were analyzed by quantitative real-time PCR in duplicate using the SYBR Green master mix (Promega, A6002, lot no. 0000472956) using primers given in **Supplementary Table S7**. Normalization was performed to input using the formula: [2^(Ct input – Ct immunoprecipitation (IP))] ^6^. Resultant values were further normalized relative to the rabbit IgG control IP values for the primer set. The relative values are represented as mean ± SD of triplicates.

**Chromatin immunoprecipitation**

The direct binding of transcription factors (HIF1α and CTCF) and RNA Pol II on target DNA sequence was evaluated by performing the ChIP-qPCR assay, as described previously ^4,5^. Briefly, cells were fixed with 1% formaldehyde and quenched by 0.125M glycine followed by cell lysis. Further, cross-linked chromatin was sonicated to an approximate chromatin fragment length of ~100–400 bp and chromatins were immunoprecipitated with HIF1α or CTCF or RNA Pol II antibody or corresponding control (immunoglobulin G) antibody (antibody details in **Supplementary Table S5**) followed by addition of ProteinG dynabeads (Invtrogen, Cat No. 10004D). The obtained complex and input DNA were reverse cross-linked in TE buffer with 1% SDS and proteinase K (Invitrogen, 25530049, lot no. 2291039 for overnight) at 65^0^C. Eluted DNA was then purified using PCR purification kit (QIAGEN, 28106). The immunoprecipitated (IP) DNA and 5% input were analyzed by qRT-PCR using specific primers (**Supplementary Table S7**) and SYBR Green Master Mix (Promega, A6002, lot no. 0000472956). All the ChIP experiments were performed at least thrice. IP DNA values were normalized to input using the following formula: 2^(Ct_input − Ct_immunoprecipitation (IP)) ^6^. Resultant values were subsequently normalized to IgG control IP DNA values. The relative values are represented as mean ± SD of triplicates.

**Chromosome Conformation Capture (3C) assay**

The 3C analysis was performed as described before ^7,8^ with slight modifications. In brief, 5 x 10^6^ MCF7 cells were fixed with 1% formaldehyde for 10 min at room temperature, and the reaction was quenched by 0.125 M glycine for 5 min at room temperature. Cross-linked cells were washed with ice cold PBS, scraped and collected. Cell pellet was re-suspended with 5ml lysis buffer (50 mM Tris-HCl with pH 8.0, 150 mM NaCl, 5 mM EDTA and 0.5% NP-40 with 1× proteinase inhibitor), and incubated on ice for 20 min. Cell pellet was then resuspended in 440ul of Milli-Q water and 60 µl 10×RE buffer (NEB, B7002S) was added, then incubated with 15 µl 10% SDS for 90 min at 37^0^C with shaking at 900 rpm. Further, 75 µl of 20% triton-X-100 was added and incubated for 90 min at 37^0^C with shaking at 900 rpm. After heating, 5 µl of aliquot was taken as undigested control. Restriction enzyme was then added sequentially 200 U HindIII (NEB, R3104T, lot No. 10029681) for 4 h at 37^0^C with shaking at 900 rpm. Again 200 U RE was added and incubated for overnight at 37^0^C with shaking. At last, final 200U RE was added and incubated for 4 h at 37^0^C with shaking. 5 µl of aliquot was taken as digested control. After inactivating the restriction enzyme for 15 min at 65^0^C, fragments were ligated with T4 DNA ligase (NEB, M0202S, lot no. 10113858) overnight at 16^0^C. Reverse crosslinking was carried out at 65^0^C for overnight with proteinase K (Invitrogen, 25530049, lot no. 2291039), followed by RNase A (Invitrogen, 12091021) treatment and phenol/chloroform extraction precipitation. The 3C interactions were analyzed by quantitative real-time PCR and product was then run on 2% agarose gel. The amount of DNA in the qPCR reactions was normalized across 3C libraries using internal control primers. Primer sequences are listed in **Supplemental Table S8**.

**Transwell invasion assays**

The transwell filter inserts (corning, 3422) were coated with matrigel (corning, 356230, lot no. 2010001) and allow to set for 3 h at 37^0^C incubator. A total of 3x10^4^ cells suspended in serum free media were placed in the upper chamber of the transwell setup and cell culture medium containing 10% FBS was added in the lower part of the well and incubated for 48-72 h in normoxic or hypoxic conditions. Non-invasive cells were gently removed from the top of the matrigel and the membrane was fixed in 4% formaldehyde followed by staining with crystal violet (0.05% crystal violet in 10% methanol in 1× PBS). Five random fields were counted using an inverted microscope (Olympus CKX41). All the invasion experiments were performed at least thrice. The % cell invaded is represented as mean ± SD of triplicates.

**Western blot**

Cells were washed with 1×PBS and protein was extracted by adding Urea lysis buffer (8 M urea, 2 M thiourea, 2% CHAPS, 1% DTT) supplemented with 1X Protease inhibitor (leupeptin 10μM, pepstatin 5 μM, EDTA 1mM, AEBSF 200 μM) and spun at 14,000 × g in 4^0^C centrifuge. Protein concentration was determined and equal amounts of protein extract were separated by SDS/PAGE, electroblotted onto PVDF membranes, and were incubated with primary antibodies followed by secondary antibody (details of antibody given in **Supplementary Table S5**). Protein bands were visualized using Odyssey Infrared imaging system (Licor). Quantification of the bands was done using ImageJ software.

**RNA isolation and cDNA synthesis**

Total mRNA was extracted using TRIzol (Invitrogen, 15596026) according to the manufacturer’s instructions. The RNA concentrations were measured using NanoDrop (Thermo Fisher Scientific, ND8000). cDNA was synthesized from 1 μg of total RNA by PrimeScript 1^st^ strand cDNA Synthesis Kit (TaKaRa, 6110A, lot no. AJX1015N) as per manufacturer’s instructions.

**Quantitative reverse transcription real-time PCR (qRT-PCR)**

Amplification reactions were performed on light cycler 480 II (Roche) using Go tag SYBR Green master mix (Promega, 75665). The gene expression were calculated using the following formula: 2^(Ct_control − Ct_target) where RPS16 is taken as control ^6^. Also, to control gene normalization, exon-level expressions were normalized to a constitutive exon. Each primer sequence is indicated in **Supplementary Table S9**.

**RNA interference**

MCF7 or HCC1806 cells (2×10^5^) were seeded in six-well culture plates. After 24 h, cells were infected with lentivirus containing small hairpin RNA (shRNA) (Sigma, Mission Human Genome shRNA Library, **Supplementary Table S10**) against *HIF1A*, *CTCF*, *DNMT1*, *DNMT3A*, *DNMT3B* or eGFP (shControl) with 8 μg/ml polybrene (Sigma, H9268) containing media. Cells were selected using 1 μg/ml puromycin (Sigma, P9620) for 3 days. Post-selection, cells were used for downstream experiments. For rescue experiments, overexpression of CTCF in *HIF1A* Knockdown cells or COL5A1exon64A in *CTCF* Knockdown cells was done 2 days post-selection using Lipofectamine 2000 reagent (Invitrogen, 11668019) as per the manufacturer’s instructions. For manipulation of HIF-1α expression in normoxic cells, we transfect *DNMT3A* Knockdown cells with a plasmid constitutively expressing a stable form of HIF-1α (HA-HIF1alpha P402A/P564A-pBabe-puro), which was a gift from William Kaelin (Addgene plasmid # 19005; http://n2t.net/addgene:19005; RRID: Addgene_19005, ^9^).

**sgRNA designing and cloning in dCas9-DNMT3A expression vector and cell transfections**

Guide sequences targeting the loci of interest were selected using the GT-Scan web-tool (<http://gt-scan.braembl.org.au/gt-scan/>, ^10^) and CRISPick (https://portals.broadinstitute. org/ gppx/crispick/public ^11^). For non-targeting (NT) control guide RNAs (sgcontrol), control plasmid (Addgene #71830) was used. All sgRNA were synthesized by oligo annealing and cloned into dCas9_DNMT3A (Addgene #74407) expression vectors via BbsI sites as described previously ^12^ using BpiI/BbsI (Fermentas, Cat No. ER1012). The cloned guide RNAs were verified by sequencing using the U6–F primer. Sequences of all oligonucleotides are listed in **Supplementary Table S11**.

Transfections were carried out using 1:3 ratio of DNA to TurfoFect (Thermo scientific, Cst No. R0531) following manufacturer's instructions. Briefly, MCF-7 cells were seeded at 80% confluency in either 60-mM (In-vitro Technologies, catalog no. FAL353046) or 15-cm culture dishes (In-vitro Technologies, catalog no. COR430599). Twenty-four hours post seeding, 3 µg of dCas9-DNMT3A constructs co–expressing dCas9-DNMT3A and a chimeric sgRNA was use to transfect cells in 60 mM dish. This was scaled up for 15-cm plate transfections. Cells were then given hypoxia treatment for 24 h. Cells were subsequently used to harvest either RNA or protein and used for other experiments such as ChIP, MeDIP, 3C and invasion assay.

**Bioinformatics analysis**

TCGA breast cancer data was analysed for CTCF and COL5A1 expression levels in normal vs tumor samples using online data portal GEPIA2.0 (<http://gepia2.cancer-pku.cn/>, ^13^). The Pearson correlation test was carried out to compare expression of CTCF gene with the hypoxia signature genes (list from MSigDB, ^14^), based on mRNA levels from TCGA BRCA dataset using online data portal GEPIA2.0 ^13^. To validate the same, further TCGA tumor samples were stratified according to CTCF expression, as described previously ^15^. Gene set enrichment analysis was performed using GSEA software to check the enrichment of hypoxia pathway genes (using the SHI_ET_AL gene signature) in CTCF-high vs CTCF_low tumor samples ^16^.

The microarray dataset in Gene Expression Omnibus (GEO) under accession number GSE147516 was used to analyze the COL5A1 gene expression and alternative splicing event for COL5A1 Exon 64 in normoxia vs hypoxia HCC1806 cell line. For HTA 2.0 triple-negative breast cancer patient profile (GSE76250) ^17^, was analysed for COL5A1 expression and COL5A1 exon 64 AS event as before ^2,15^. Splicing events with absolute splicing index (|SI|) ≥1.5 and P < 0.05 were considered to be significant.

**In Silico analysis for CpG Islands and transcription factor binding sites in *CTCF* and *COL5A1* promoter region**

Human *CTCF* and *COL5A1* gene promoter sequence was retrieved from the Eukaryotic Promoter Database (<https://epd.vital-it.ch/>, ^3^). The CpG islands within 2000 bp upstream and 500 bp downstream of *CTCF* and *COL5A1* gene TSS start site (transcript NM_006565 and NM_000093, respectively) were identified using DBCAT (<http://dbcat.cgm.ntu.edu.tw/>, ^18^) and EMBOSS CpGPlot (<https://www.ebi.ac.uk/emboss/cpgplot>, ^19^). The putative transcription factor binding sites were predicted using JASPAR ^20,21^ and CIS-BP ^22^.

**Retrieval of gene list from various databases and overlapping of data set to found CTCF novel targets**

List of genes significantly upregulated in breast cancer tissue in comparison to the normal tissue was obtained from GEPIA2. List of genes hallmark of EMT and hypoxia were obtained from MSigDB. Gene overlapping from different sets was performed using online tool <https://bioinformatics.psb.ugent.be/webtools/Venn/>.

**In silico analysis of CTCF Binding motifs for the *COL5A1* locus**

We used Find Individual Motif Occurrences (FIMO) software ^23^ to search CTCF Binding motifs within the COL5A1 gene with default parameters and *p*< 0.0001 or 0.01. The binding motif for CTCF (Matrix ID MA0139.1) from the JASPAR database ^21^ was taken as inputs. The program generates both the forward and reverse strand hits which are ranked to a logarithmic sequence similarity score on binding locations.

**RNA-seq data analysis**

MCF7 cells normoxia and hypoxia (16 h) RNA-seq data was obtained from GSE166203. Gene expression levels and differentially expressed genes (DEGs) in normoxia vs hypoxia (16 h) were identified using NOIseq analysis ^24^. Genes with probability of difference of 0.6 or more were called as differentially expressed genes. Alternative splicing events in hypoxia as compared to the normoxia were retrived using rMATS (v. 4.0.2) ^25^.

**ChIP-Sequencing**

ChIP experiment was performed as above. ChIP-seq experiments were carried out at Core Technologies Research Initiative (CoTeRI), National Institute of Biomedical Genomics (NIBMG, University of Kalyani, Kolkata, West Bengal, India).  ChIP-seq library preparation was performed using TruSeqChIP Sample Prep Kit (Illumina) according to the manufacturer's instructions. 10 ng of input ChIP-enriched DNA was used for ChIP-seq library preparation. Paired-end sequencing (2 × 100 bp) of these libraries were performed in Novaseq 6000 platform.

**ChIP-seq data analyses**

For each library, raw fastq files were trimmed by Trimmomatic (v0.39) with default parameters ^26^. Reads were uniquely aligned to GRCh38 using STAR (v2.7.3a) aligner ^27^. The biological replicates were merged by Samtools v1.9 ^28^. MACS2 v2.1.2 was used to call peaks with default parameters ^29^. To identify the CTCF differential binding regions in normoxia vs hypoxia CTCF ChIP-seq, DiffBind v3.6.5 was used ^30^. The peaks called by MACS2 and differential peaks identified by DiffBind were annotated by using BioMart by ENSEMBL ^31^. The GO analysis was performed using ShinyGO (v0.76.2) ^32^ and Enrichr ^33^. Integrative Genomics Viewer was use to visualize the signals ^34^.

To find CTCF mediated exon inclusion events under hypoxia, we quantified the distribution of hypoxia-gained CTCF sites with exons included under hypoxia using BEDtools and were put into three groups i) CTCF peak either overlapping with exon or intronic regions downstream of exons corresponding the roadblock model discussed above ^5^ ii) CTCF peak present at promoter ±15 kb and also in intronic regions upstream of exons reflecting CTCF-mediated promoter-intragenic looping upstream of exon ^35^ iii) CTCF peak present at promoter ±15 kb and also in intronic regions both upstream and downstream of exons correlating present study mechanism coupling promoter-exon upstream looping with CTCF mediated RNA Pol II pause at exon.

**Supplementary Table S4: Clinical characteristics of patients.**

| **S. No.** | **Patient No.** | **Histopathology** |
| --- | --- | --- |
| 1 | Patient 1 | Infiltrating duct carcinoma ypT4N0Mx |
| 2 | Patient 2 | Infiltrating duct carcinoma pT2NMx |
| 3 | Patient 3 | Invasive carcinoma of no special type duct grade II |
| 4 | Patient 4 | Invasive carcinoma of no special type grade II pT2N1Mx |
| 5 | Patient 5 | Infiltrating duct carcinoma Grade III |
| 6 | Patient 6 | Infiltrating duct carcinoma Grade II |
| 7 | Patient 7 | Infiltrating duct carcinoma stage pT2N2Mx |
| 8 | Patient 8 | Infiltrating duct carcinoma Grade II pT3N0Mx |
| 9 | Patient 9 | Mucinous carcinoma pT2SnNMx |
| 10 | Patient 10 | Invasive carcinoma of no special type ductal grade II pT2(Sn)N0mx |
| 11 | Patient 11 | Invasive carcinoma of no special type grade II pT2NxMx |
| 12 | Patient 12 | Invasive carcinoma of no special type grade II pT3N1Mx |
| 13 | Patient 13 | Invasive carcinoma of no special type ductal grade III pT2N2Mx |
| 14 | Patient 14 | Infiltrating duct carcinoma Grade III pT4N0Mx |
| 15 | Patient 15 | Invasive carcinoma of no special type ductal grade II pT2N1Mx |
| 16 | Patient 16 | Invasive carcinoma of no special type ductal grade II pT2N1Mx |
| 17 | Patient 17 | Infiltrating duct carcinoma pT2N2Mx |
| 18 | Patient 18 | Infiltrating duct carcinoma Grade II pT2N0Mx |

**Supplementary Table S5: List of antibodies used ChIP, MeDIP and western blot assays**

| **S.No** | **Antibody** | **Company** | **Catalog no** | **Lot no** |
| --- | --- | --- | --- | --- |
| 1 | CTCF (D31H2) | CST | 3418S | 1 |
| 2 | HIF-1α (D2U3T) | CST | 14179S | 3 |
| 3 | DNMT1 (D63A6) | CST | 5032S | 3 |
| 4 | DNMT3A (D2H4B) | CST | 32578S | 1 |
| 5 | DNMT3B (E8a8a)xp | CST | 57868S | 1 |
| 6 | GAPDH (D16h11) | CST | 5174S | 8 |
| 7 | COL5A1 | CST | 37304S | 1 |
| 8 | Snail (C15D3) | CST | 3879S | 12 |
| 9 | Anit-Flag tag | Novus Biologicals | NBP1-06712SS | B-6 |
| 10 | Anti-E cadherin | Abcam | Ab40772 | GR148899-1 |
| 11 | Anti-Vimentin | Abcam | ab137321 | GR294886-5 |
| 12 | Alexa-Flour 680 anti-rabbit IgG | Invitrogen | A32734 | RJ243414 |
| 13 | Alexa-Flour 800 anti-mouse IgG | Invitrogen | A32730 | SC243837 |
| 14 | Anti-Carbonic Anhydrase | Abcam | ab184006 | GR173128-25 |
| 15 | Alexa-Flour 680 anti-rat IgG | Invitrogen | A21096 | 2010149 |
| 16 | Cas9 (7A9-3A3) | CST | 14697S | 3 |
| 17 | H3K4trimethylation (H3K4me3) | Abcam | Ab8580 | GR273043-6 |
| 18 | H3K27acetylation (H3K27ac) | Abcam | Ab4729 | GR312651-3 |
| 19 | Rpb1 CTD (4H8) (RNA Polymerase II CTD) | CST | 2629S | 3 |
| 20 | 5-methylCytosine (D3s2z) | CST | 28692S | 2 |
| 21 | Normal Rabbit IgG | CST | 2729S | 10 |
| 22 | Normal Mouse IgG | Millipore | NI03-100UG | 3129279 |

**Supplementary Table S6: List of primers used for molecular cloning**

| **CTCF Luciferase construct** | **Forward primer 5'-3'** | **Reverse primer 5'-3'** |
| --- | --- | --- |
| CTCF (+687 to -145) | GCTTTTAGGTGGCCCTCTTC | CTTACCTCCGCGCCACAC |
| CTCF HRE Mutant | GGCGCGTGCGTTTTGACGACGCGAC | CTCGCCGGGTCGCTCGTGGGC |
| **Over expression full length construct** |  | |
| CTCF | ATGGAAGGTGATGCAGTCGAAG | TCACCGGTCCATCATGCTGAGG |
| COL5A1 exon 64A isoform | ATGGACGTCCATACCCGCTGGA | CTAGCCCATGAAGCAAGCCGGCCC |

**Supplementary Table S7: List of primers used for ChIP/MeDIP-qPCR**

| **ChIP/MeDIP-qPCR** | **Forward primer 5'-3'** | **Reverse primer 5'-3'** |
| --- | --- | --- |
| CTCF promoter (+320 to +157 from TSS) | ACCCAGCGGGAGCCAGGAC | GTTGGCGCAGGGCAGCAT |
| COL5A1 promoter (+54 to -47 from TSS) | GAGAGAGGGAGGGAGGAAAA | CTCCGCTCGAGTGAGGTC |
| COL5A1Exon 64A | GTTTCTCTCCCTCCCCACCT | CCAGAGGCCACCTTACCAG |
| COL5A1Exon 64B | CCGCCCATCTCGTATCTTAC | CGAGGTGGCCACTTACCAG |
| COL5A1_T1 | GACAAGAGCAGTTAGCTGCCCT | AGCAGGGGCCAGCCCTCAGC |
| COL5A1_T2_1 | CCAAGGCTGCAAACTGTTAAG | CCATCCCTTCCTCCAGTCTC |
| COL5A1_T2_2 | GCCAATCCAGGCATCCAG | CCGAAGATCTCTTCCATGC |
| COL5A1_T3 | GGACACATTGATGTGAGGACA | TTGCTGGGAGAAAAGACAGAA |

**Supplementary Table S8: List of 3C Assay Primers**

| **3C-qPCR** | **Forward primer 5'-3'** | **Reverse primer 5'-3'** |
| --- | --- | --- |
| Internal control | CTCCTCCTTTCTGATGCCTCTGTTCAC | CTCTGTCCCTAAGAGGTGTTCCTCTCGT |
| COL5A1_C | ATTACTGAACTCCAGGCCAGCAAGGT | GGAACACTGCTCTTTCCTCCTGCAAAAC |
| COL5A1_T1 | CTAGCTCCGAGGGAATTGAGAAAGCAAC | CAGGATGCACGTACACATACAGACATGC |
| COL5A1_T2 | GGTCAGGAAGGGTTGGAGGGCTTTC | CCCACCCCGAATTCTTCACTAAACAGAC |
| COL5A1_T3 | CACTCAACACATACAAACCTGGCAGACC | TTGTCACAGTGTCTGTCTTCCCTAGCAG |

**Supplementary Table S9: List of primers used for gene expression analysis**

| **RT-PCR** | **Forward primer 5'-3'** | **Reverse primer 5'-3'** |
| --- | --- | --- |
| CTCF exon 5 | ATGTTATATTTGTCATGCTCGGTTTA | TTTCGGGCTATGACTGTGTC |
| COL5A1 exon2 (constitutive exon) | AGCAGATCTCCTGAAGGTTCT | GGGTACAGCTGCTTGGTG |
| COL5A1 exon 64A | CCGAAGGGGCCAGAATCACT | CTCGGCGTCCACATAGGAGA |
| COL5A1 exon 64B | GAAGGGAGTAAAATGGCCCGC | CTCGGCGTCCACATAGGAGA |
| RPS16 | AAACGCGGCAATGGTCTCATCAAG | TGGAGATGGACTGACGGATAGCAT |

**Supplementary Table S10: List of shRNA used for gene knockdown**

| **shRNA** | **Target Sequence 5'-3'** |
| --- | --- |
| shCTCF_1 | GTACCGGTATGATTTCCCATCGACATTTCTCGAGAAATGTCGATGGGAAATCATATTTTTTG |
| shCTCF_2 | CCGGGCGGAAAGTGAACCCATGATACTCGAGTATCATGGGTTCACTTTCCGCTTTTT |
| shHIF1A_1 | CCGGGTGATGAAAGAATTACCGAATCTCGAGATTCGGTAATTCTTTCATCACTTTTT |
| shHIF1A_2 | CCGGTGCTCTTTGTGGTTGGATCTACTCGAGTAGATCCAACCACAAAGAGCATTTTT |
| shDNMT1_1 | CCGGGCCCAATGAGACTGACATCAACTCGAGTTGATGTCAGTCTCATTGGGCTTTTT |
| shDNMT1_2 | CCGGCGACTACATCAAAGGCAGCAACTCGAGTTGCTGCCTTTGATGTAGTCGTTTTT |
| shDNMT3A_1 | CCGGCCGGCTCTTCTTTGAGTTCTACTCGAGTAGAACTCAAAGAAGAGCCGGTTTTTG |
| shDNMT3A_2 | CCGGGCCTCAGAGCTATTACCCAATCTCGAGATTGGGTAATAGCTCTGAGGCTTTTTG |
| shDNMT3B_1 | CCGGGCAGGCAGTAGGAAATTAGAACTCGAGTTCTAATTTCCTACTGCCTGCTTTTTG |
| shDNMT3B_2 | CCGGCCATGCAACGATCTCTCAAATCTCGAGATTTGAGAGATCGTTGCATGGTTTTTG |
| shControl | 5’-CCGGTACAACAGCCACAACGTCTATCTCGAGATAGACGTTGTGG  CTGTTGTATTTTT-3’ |

**Supplementary Table S11: List of primer sequences used to Construct Guide RNAs for dCas9-DNMT3A system**

| **dCas9-DNMT3A-sgRNA** | **Target Sequence 5'-3'** |
| --- | --- |
| CTCF-HRE-sgRNA1 | CGTGGGCTCTGCCGGCGCCG |
| CTCF-HRE-sgRNA2 | GGCAGAGCCCACGAGCGACC |
| CTCF-HRE-sgRNA3 | CGACCCGGCGAGGGCGCGTG |
| COL5A1 Exon 64A-SgRNA1 | CACCTCCCCGCTGCATGTTT |
| COL5A1 Exon 64A-SgRNA2 | TCTTGGCCCAAAGAAAACCC |
| COL5A1 Exon 64A-SgRNA3 | TGAATTCAAGCGTGGGAAAC |
| COL5A1 Exon 64A-SgRNA4 | GGTGGCCTCTGGCGTCTTTG |
| COL5A1_T1-sgRNA1 | CTGCAGGTACACATGGCCCC |
| COL5A1_T1-sgRNA2 | GGGGGCCCCTGGCACGCTGA |
| sgControl | ACGGAGGCTAAGCGTCGCAA |

**Supplementary Figure S1**

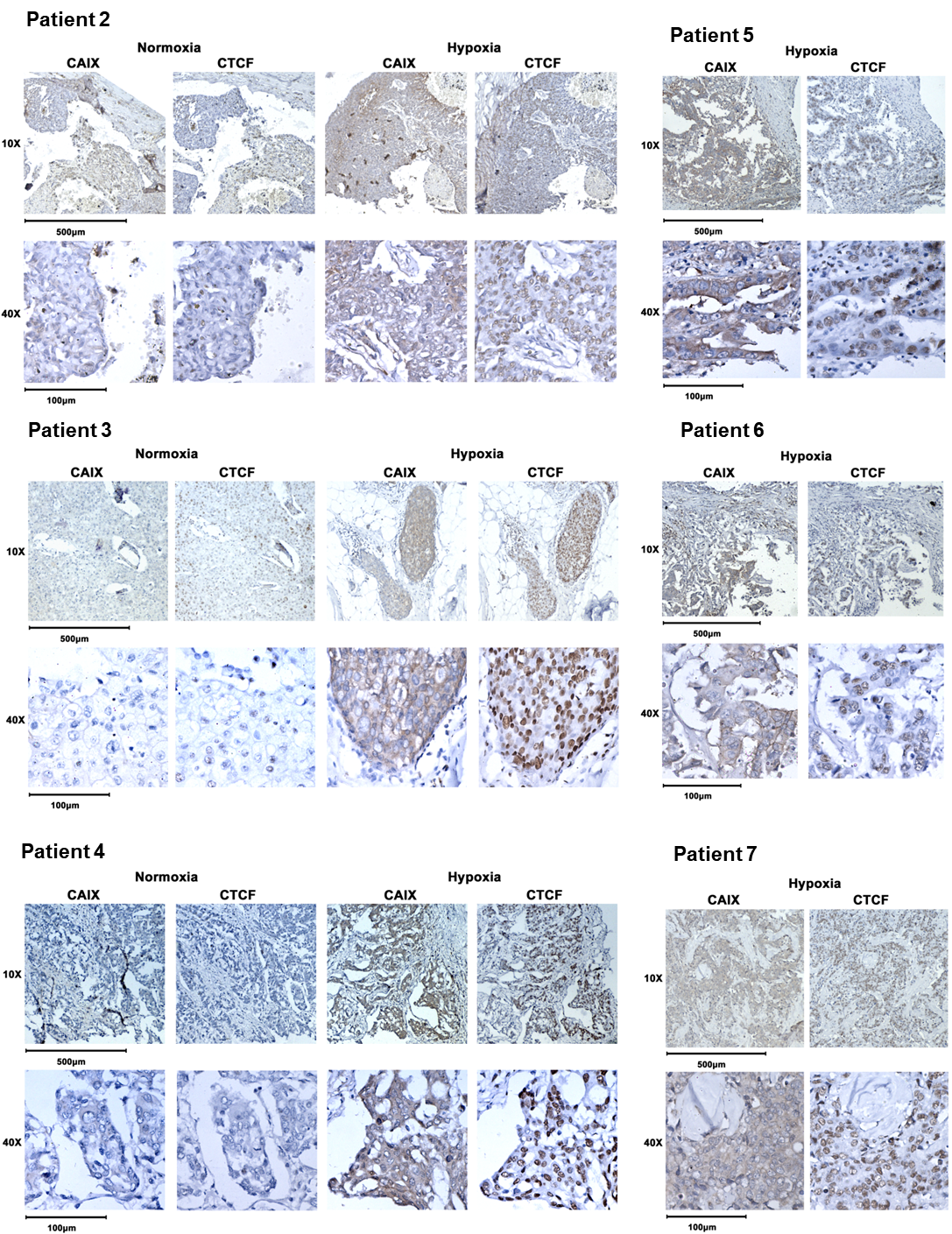

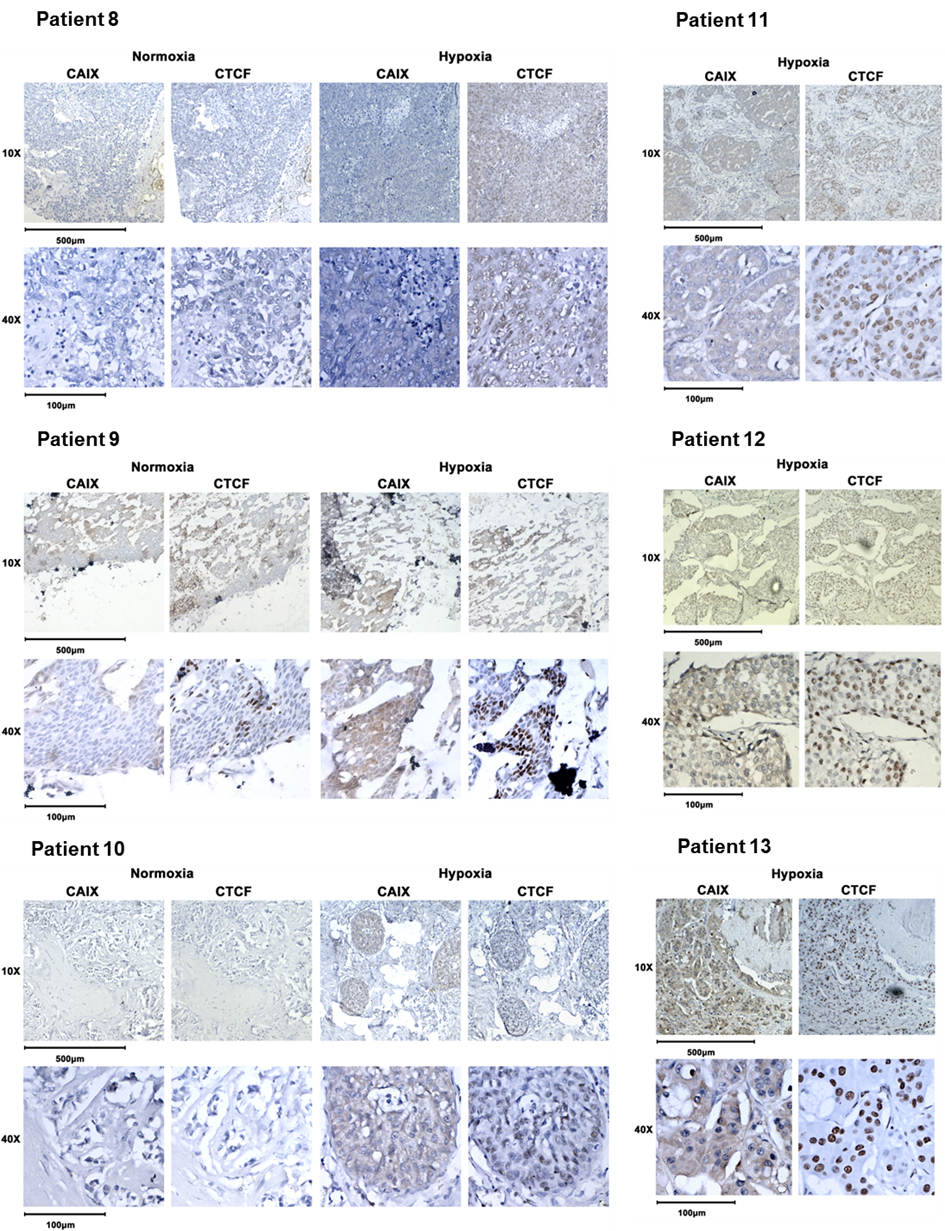

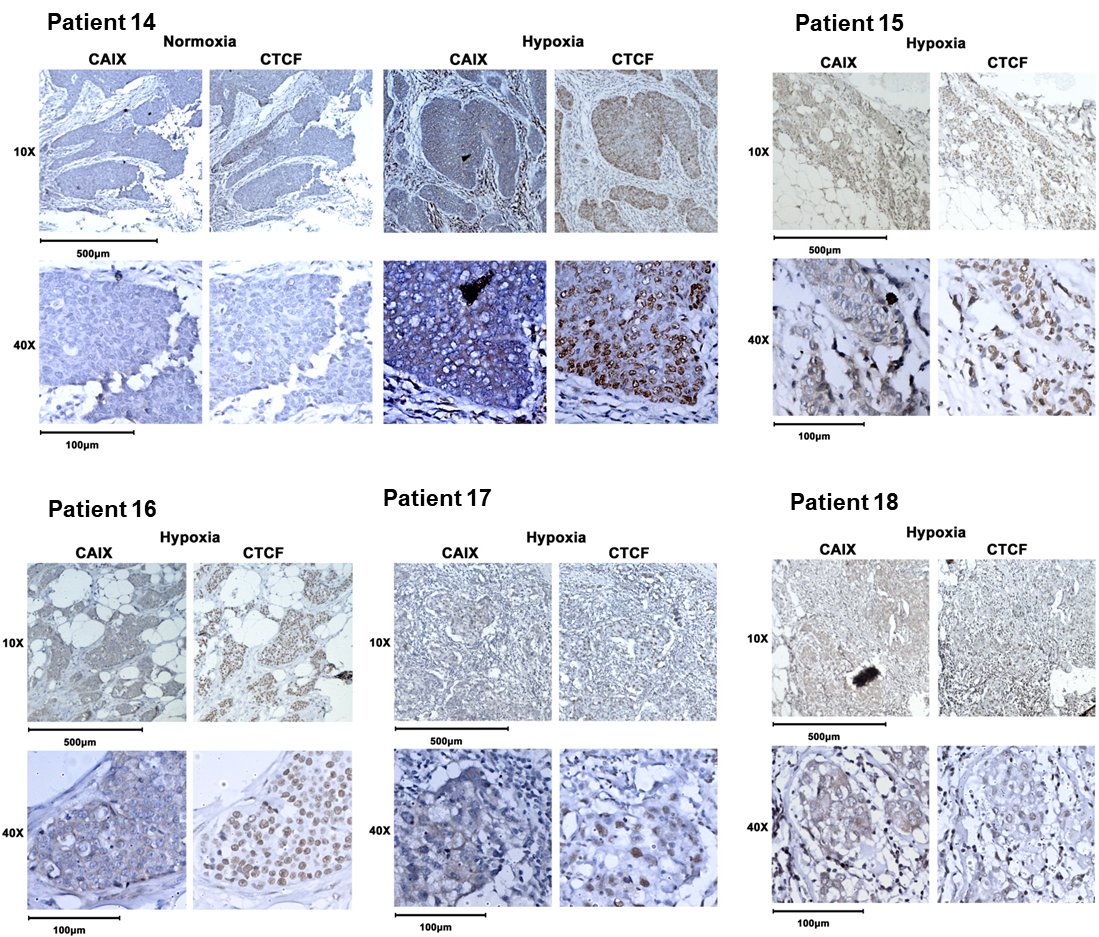

**Supplementary Figure S1:** **CAIX, and CTCF immunostaining of 17 illustrative cases of breast cancer patients.** Hypoxic regions: Areas representing strong membranous and/or cytoplasmic immunostaining for CA IX also exhibit strong expression of CTCF (nuclear) and Normoxic regions: Areas representing weak immunostaining for CAIX also exhibit weak expression of CTCF. Magnification: 10× and 40×.

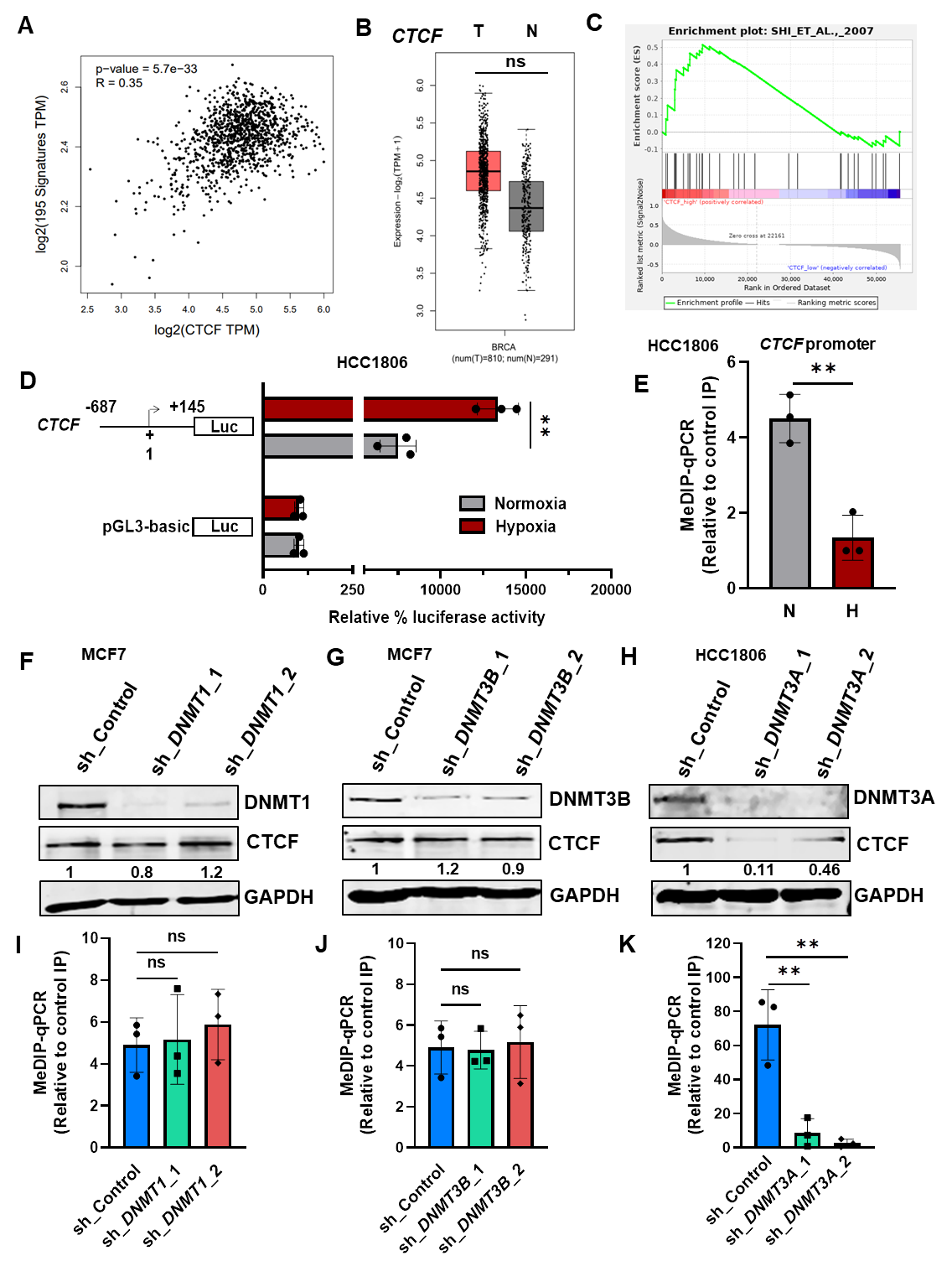
**Supplementary Figure S2**

**Supplementary Figure S2:** **CTCF is induced in hypoxic breast cancer.** **A)** The mRNA expression of *CTCF* was compared with a 195-gene hypoxia signature in 1,218 human breast cancers from TCGA database by Pearson’s correlation test **B)** TCGA gene expression profile of *CTCF* in normal breast tissue and primary breast tumor **C)** GSEA enrichment plot for hypoxia gene signature (Shi et al., 2007) for TCGA breast cancer samples stratified as *CTCF*_high (n=110) or *CTCF*_low (n=110) **D)** HCC1806 cells were co-transfected with *CTCF* promoter-luciferase construct (-687 to +145) and pRL-TK Renilla, exposed to normoxia and hypoxia. The relative luciferase values are shown as mean ± SD **E)** MeDIP-qPCR on *CTCF* promoter in normoxic vs hypoxic HCC1806 cells **F)** Immunoblot of DNMT1 and CTCF in normoxic MCF7 cells transduced with shRNA against *DNMT1* or shControl **G)** Immunoblot of DNMTB and CTCF in normoxic MCF7 cells transduced with shRNA against *DNMT3B* or shControl **H)** Immunoblot of CTCF and DNMT3A in normoxic HCC1806 cells transduced with shRNA against *DNMT3A* and or shControl **I)** MeDIP-qPCR on *CTCF* promoter in normoxic MCF7 cells transduced with shRNA against *DNMT1* versus shControl **J)** MeDIP-qPCR on CTCF promoter in normoxic MCF7 cells transduced with shRNA against *DNMT3B* versus shControl **K)** MeDIP-qPCR on CTCF promoter in normoxic HCC1806 cells transduced with shRNA against *DNMT3A* versus shControl.

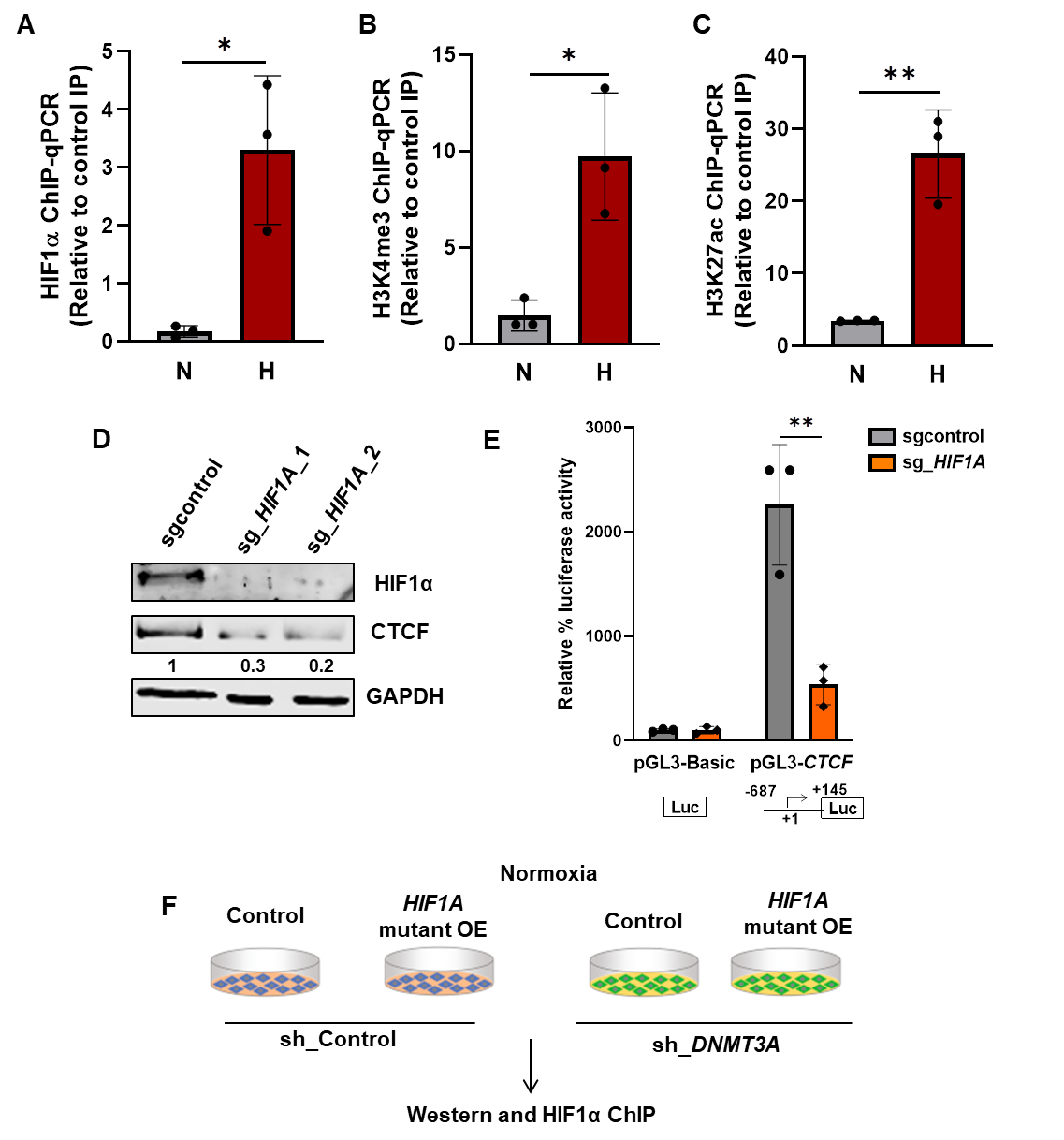
**Supplementary Figure S3**

**Supplementary Figure S3: *CTCF* is a direct target of HIF1α**. **A)** HIF1α ChIP-qPCR **B)** H3K4me3 ChIP-qPCR **C)** H3K27ac ChIP-qPCR on *CTCF* promoter in normoxic and hypoxic HCC1806 cells **D)** Immunoblot of CTCF after *HIF1A* knockout in HCC1806 cells under hypoxia **E)** Relative luciferase activity of *CTCF* promoter in HCC1806 cells transfected with sgRNA against *HIF1A* vs sgcontrol under hypoxia.

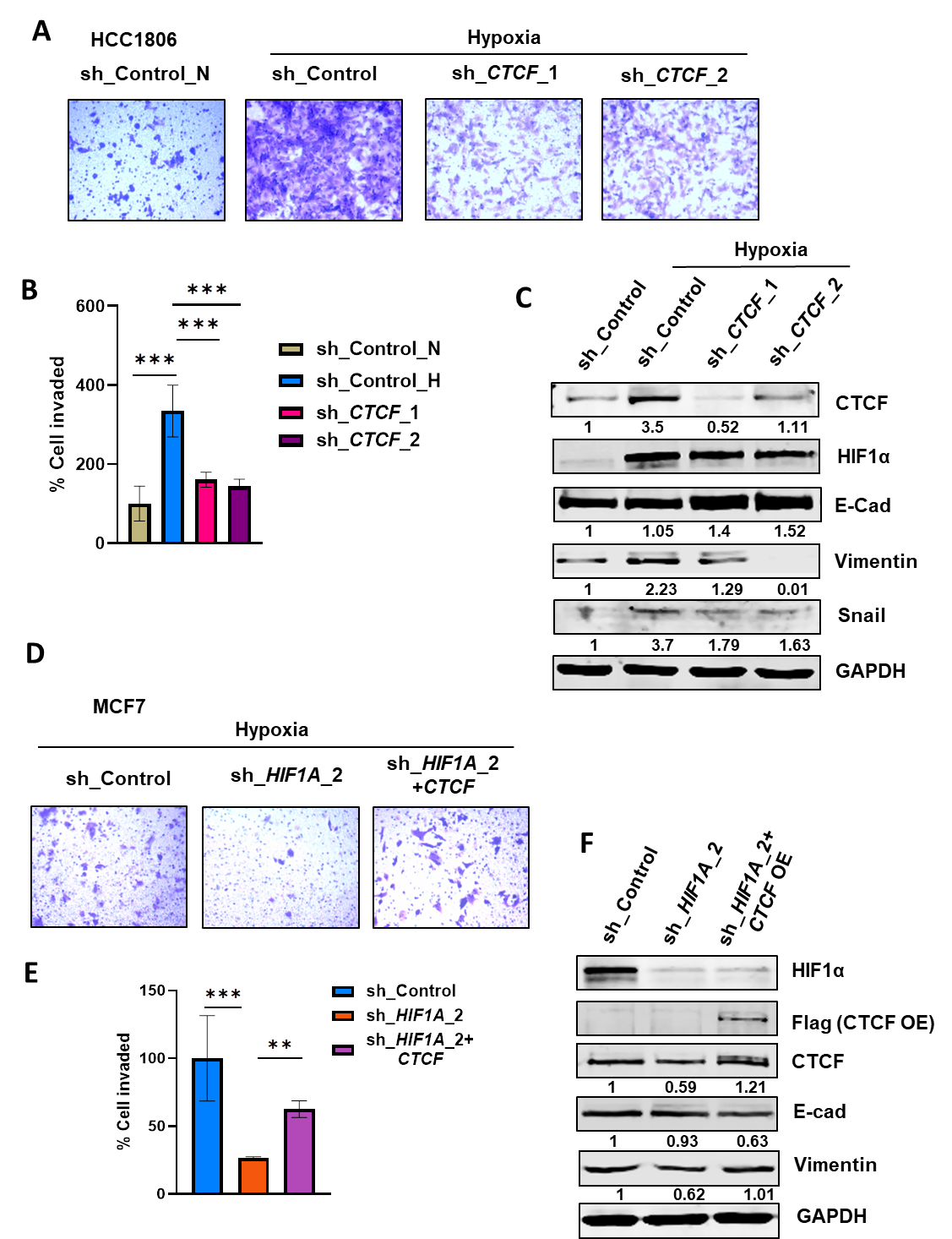
**Supplementary Figure S4**

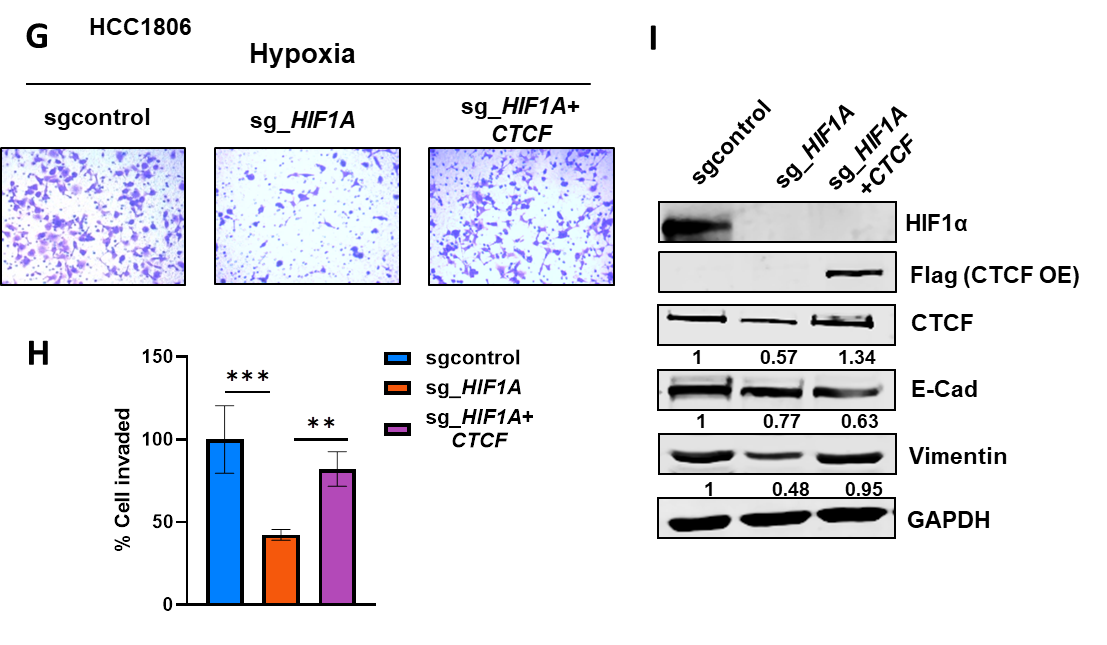

**Supplementary Figure S4: Hypoxia induced HIF1α-CTCF signaling axis promotes breast cancer cell invasive phenotype**. **A)** Invasion assay with **B)** its quantification as % of cells invaded after *CTCF* knockdown in comparison to the control HCC1806 cells under normoxic and hypoxic condition **C)** Immunoblot of CTCF, HIF1α, E-cadherin (E-Cad), vimentin and snail in *CTCF* knockdown or control HCC1806 cells under normoxic and hypoxic condition **D)** Invasion assay with **E)** quantification as % of cells invaded after overexpression of empty vector (EV) or *CTCF* in cells transduced with sh*HIF1A*_2 in comparison to the shcontrol under hypoxia in MCF7 **F)** Immunoblot of HIF1α, Flag (to confirm overexpression of CTCF), CTCF, E-cad and vimentin in shcontrol versus sh*HIF1A*_2 cells with ectopically expressed either empty vector (EV) or *CTCF* in MCF7 hypoxic cells **G)** Invasion assay with **H)** quantification as % of cells invaded after overexpression of empty vector (EV) or *CTCF* in cells transduced with sg*HIF1A* in comparison to the sgcontrol under hypoxia in HCC1806 **I)** Immunoblot of HIF1α, Flag (to confirm overexpression of CTCF), CTCF, E-cad and vimentin in sgcontrol versus sg*HIF1α* cells with ectopically expressed either empty vector (EV) or *CTCF* in HCC1806 hypoxic cells.

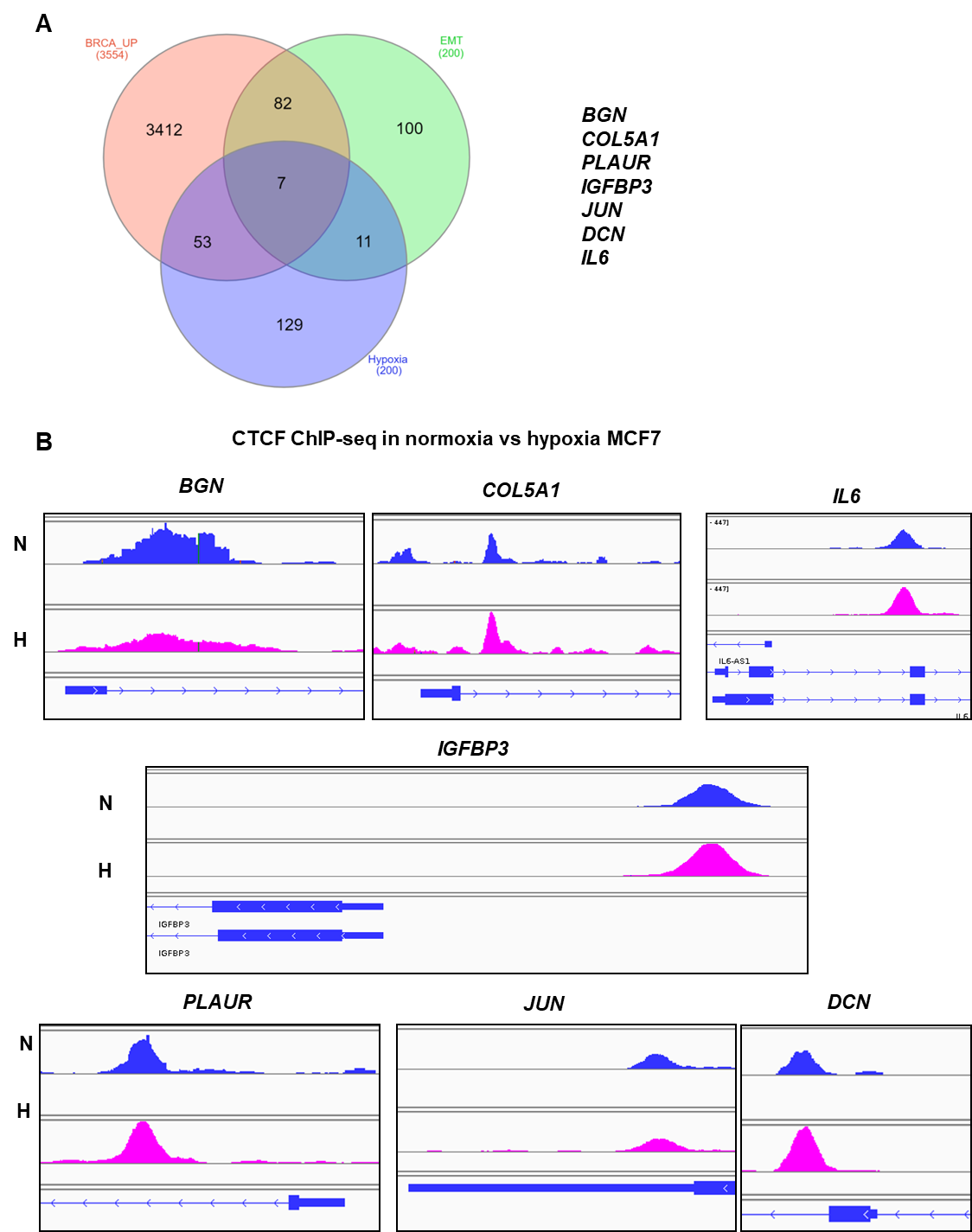
**Supplementary Figure S5**

**Supplementary Figure S5: *COL5A1* gene as a novel target of CTCF.** **A)** Venn diagram depicting the overlap and distinct genes in list of genes upregulated in breast cancer (obtained from TCGA dataset) and list of genes as a hallmark of EMT and hypoxia signature hallmark dataset (obtained from MsigDB). **B)** CTCF peak on gene promoter ±5kb in normoxia vs hypoxia condition obtained from CTCF-ChIP-seq.

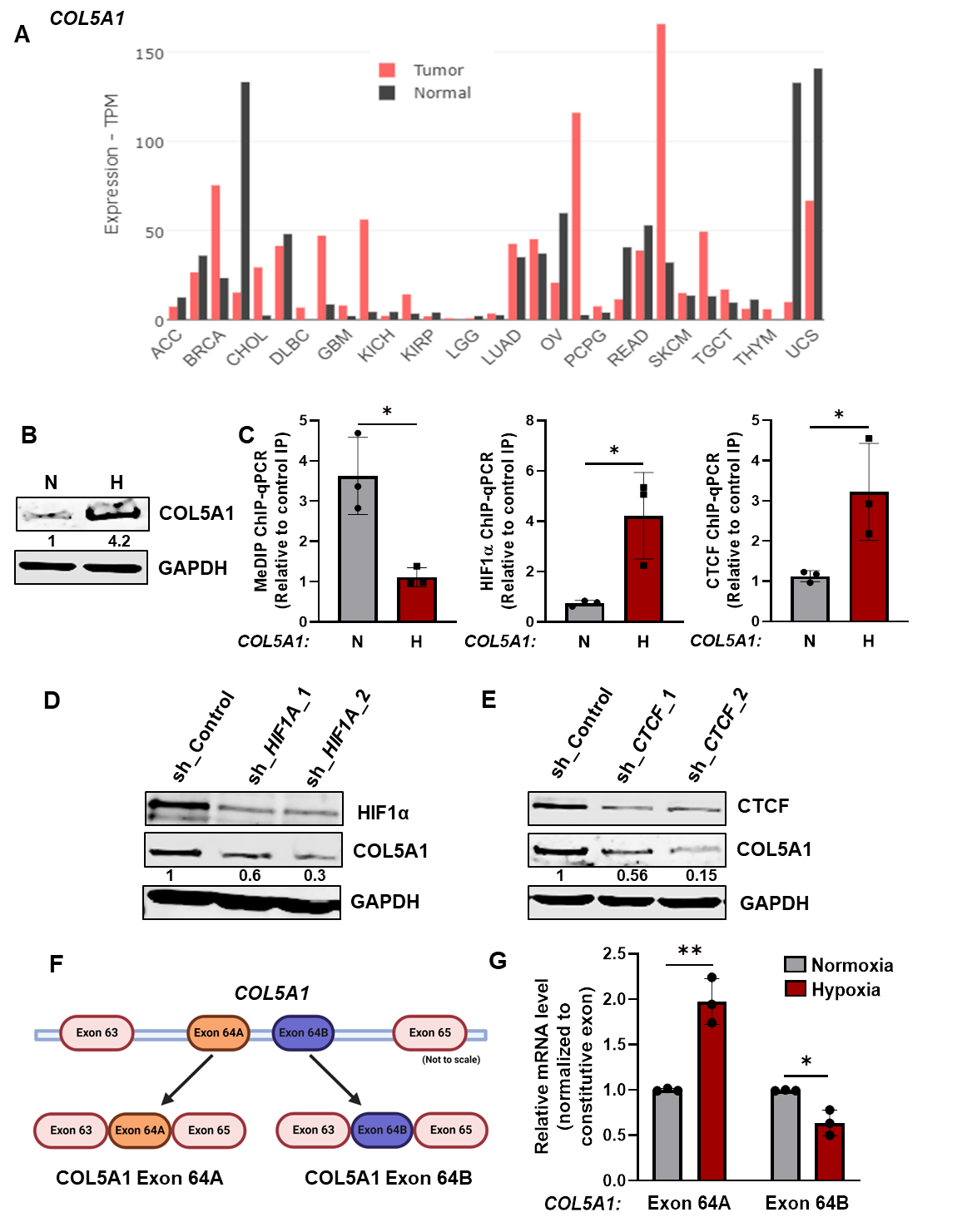
**Supplementary Figure S6**

**Supplementary Figure S6:** ***COL5A1* expression is induced under hypoxia in HIF1α-CTCF-dependent manner.** **A)** TCGA gene expression profile of *COL5A1* pertaining to normal tissue and primary tumor across different cancers **B)** Immno-blot of COL5A1 in normoxic and hypoxic breast cancer cell line, HCC1806 **C)** MeDIP, HIF1α and CTCF ChIP-qPCR on *COL5A1* promoter in normoxic and hypoxic HCC1806 cells **D)** Immunoblot of COL5A1 in HIF1α depleted HCC1806 cells under hypoxia in comparison to the shControl cells **E)** Immunoblot of COL5A1 in CTCF depleted HCC1806 cells under hypoxia in comparison to the shControl cells **F)** *COL5A1* gene exon 64 has two mutually exclusive isoforms, exon 64A and exon 64B **G)** qRT-PCR analysis of *COL5A1* exon 64A and exon 64B isoform normalized to RPS16 and constitutive exon expression levels in normoxic and hypoxic HCC1806 cells.

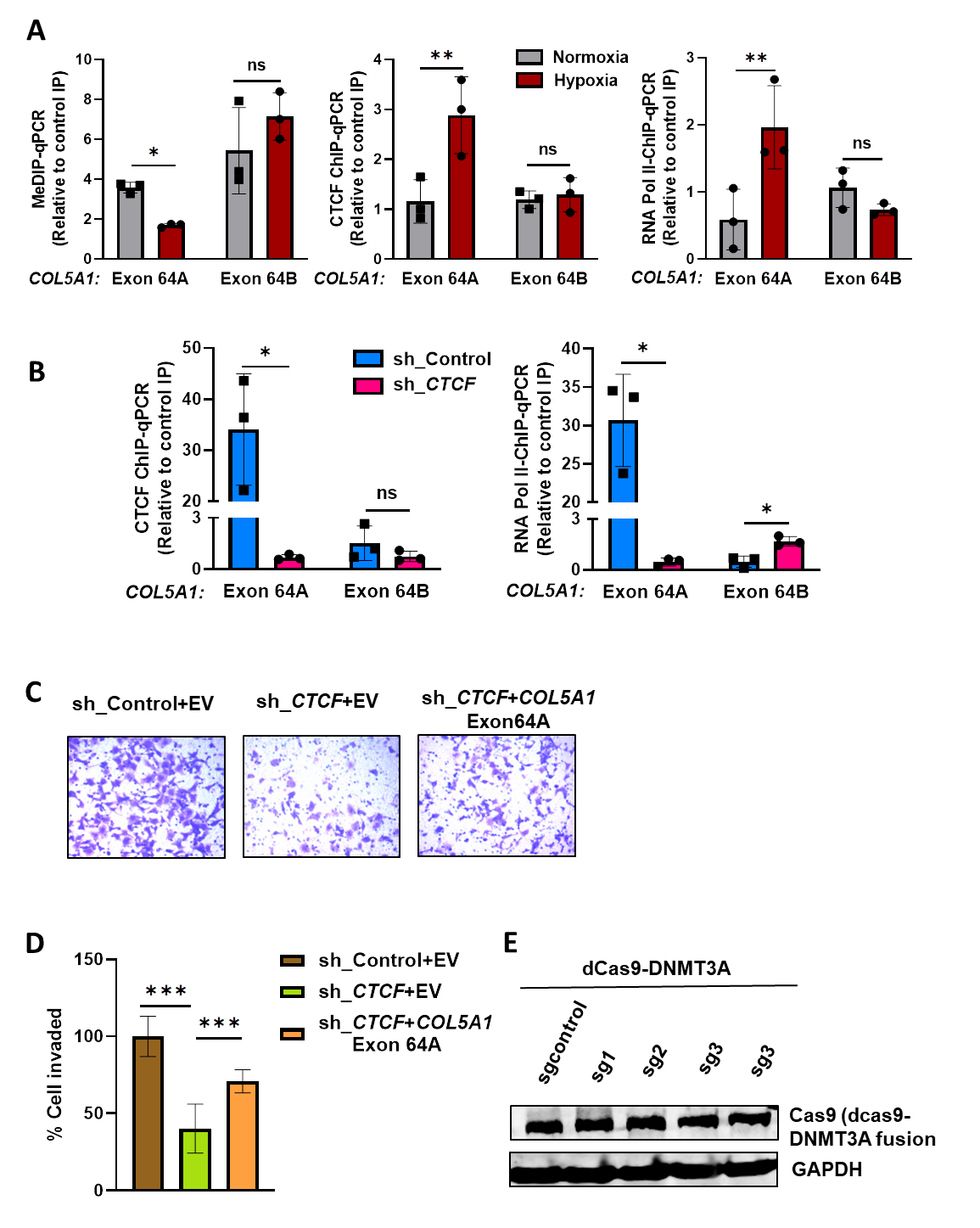
**Supplementary Figure S7**

**Supplementary Figure S7: Hypoxia-induced *COL5A1* exon 64AA inclusion is regulated by methylation dependent enrichment of CTCF on exon 64A and is associated with RNA Pol II pause. A)** MeDIP, CTCF and RNA Pol II ChIP-qPCR on exon 64A and exon 64B of *COL5A1* in normoxic and hypoxic HCC1806 cells **B)** CTCF and RNA Pol II ChIP-qPCR on exon 64A and exon 64B in HCC1806 cells transduced with either sh*CTCF* or shRNAcontrol under hypoxia **C)** Invasion assay with **D)** its quantification as % of cells invaded after over expression of either empty vector or *COL5A1*exon 64A isoform in CTCF depleted MCF7 cells in comparison to the shControl cells under hypoxia **E)** Immunoblot of Cas9 (for dCas9-DNMT3A fusion protein) in MCF7 cells transfected with dCas9-DNMT3A-sgRNAs or sgcontrol directed on exon 64A of *COL5A1* under hypoxia.

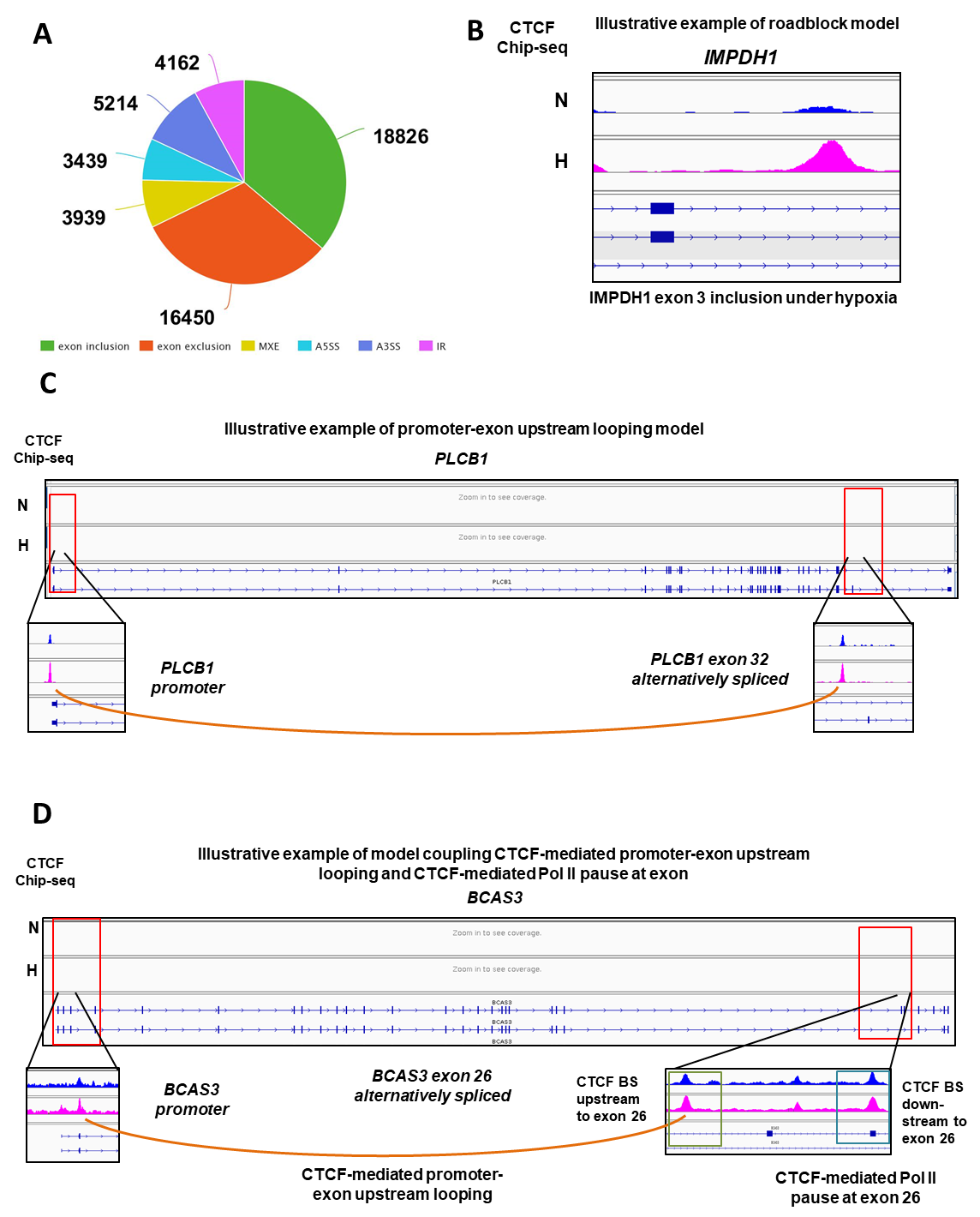
**Supplementary Figure S8**

**Supplementary Figure S8: Hypoxia-induced differential CTCF occupancy regulates transcriptional and alternative splicing dynamics in breast cancer cells to drive EMT. A)** Pie chart showing number of alternative splicing events obtained from GSE166203 using rMATS **B)** illustrative examples of exon inclusion under hypoxia by roadblock model **C)** illustrative examples of exon inclusion under hypoxia by promoter-exon upstream looping model **D)** illustrative examples of exon inclusion under hypoxia by model CTCF-mediated promoter-exon upstream looping and CTCF-mediated Pol II pause at exon showed in the present study.
